# MEM-CALVADOS: A Residue-Level Model for Flexible Proteins at Membrane Interfaces

**DOI:** 10.64898/2026.08.12.743853

**Authors:** Riccardo Saltutti, Giulio Tesei

## Abstract

Many membrane proteins contain intrinsically disordered regions (IDRs) that play key biological roles by providing structural plasticity, harboring sites for post-translational modifications, and mediating protein clustering and phase separation. Residue-level molecular models parameterized against experimental data have provided insights into how IDR sequence controls conformational properties and phase behavior in soluble proteins. Here, we extend this modeling framework to membrane-associated IDRs. We adapt a coarse-grained lipid model, iSoLFv2, for four phospholipids and combine it with CALVADOS, a residue-level model for IDRs and multi-domain proteins. Protein–lipid cross-interaction parameters are calibrated to reproduce predicted insertion and orientation of transmembrane proteins with diverse architectures. We validate the resulting model against Wimley–White free energies of transfer of hydrophobic peptides and against an experimentally refined conformational ensemble of a flexible membrane receptor. Finally, we show the applicability of the model to a membrane-associated assembly of signaling proteins. The model provides a computationally efficient framework for studying conformational ensembles and assembly of proteins at bilayer–water interfaces.

## Introduction

Transmembrane proteins are central to cellular processes such as signal transduction and molecular transport, and are targets of a large fraction of approved drugs [1]. Their membrane-spanning regions often adopt well-defined architectures, including *β*-barrels, *α*-helical bundles, and single-pass helices. In contrast, their extra- and intracellular regions are often structurally more heterogeneous and frequently contain intrinsically disordered regions (IDRs), which confer conformational flexibility [2]. IDRs in membrane proteins play important functional roles, including harboring phosphorylation sites and short linear motifs that mediate interactions with the membrane and binding partners. Changes in the lipid composition of the membrane, ligand binding, and post-translational modifications can drive conformational changes in membrane-associated IDRs, contributing to the regulation of protein function [2–6]. The same stimuli can also promote self-association of transmembrane proteins into oligomers as well as nonstoichiometric clusters [7, 8]. Lateral segregation into clusters can be coupled to the formation of biomolecular condensates, where transmembrane proteins phase separate with cytosolic or extracellular binding partners. By compartmentalizing proteins at membrane surfaces, membrane-associated condensates can increase the local concentration and dwell time of signaling proteins, thereby facilitating signal transduction [8]. The lipid membrane influences protein assembly through its composition and physical properties. In turn, membrane-associated condensates can remodel the membrane through changes in lateral lipid segregation, lipid packing, and membrane curvature [9–14].

Studying transmembrane proteins containing long IDRs poses technical challenges related to their flexibility, molecular size, and requirement for amphiphilic environments for solubilization [2, 15, 16]. Experimentally derived conformational ensembles of full-length transmembrane proteins are therefore scarce and often require integrative approaches that combine data from multiple techniques, frequently applied to separately studied protein domains [15, 16].

AlphaFold2 has greatly increased the number of available structural models for transmembrane proteins [17]. However, for transmembrane proteins containing IDRs, AlphaFold2 models can violate geometric constraints imposed by the membrane, for example by positioning disordered segments or soluble helices within the transmembrane region [18]. Recent AlphaFold3-based approaches may partly address these limitations by allowing lipid molecules to be included in the prediction [19, 20]. Even when the overall membrane topology is correctly predicted, AlphaFold models represent only a limited set of conformations and do not fully capture the large, heterogeneous conformational ensembles of IDRs, or their responsiveness to the molecular environment [21–24].

For flexible membrane proteins, conformational ensembles and their modulation by the local lipid composition and membrane properties are often strongly related to function. Molecular dynamics simulations provide a useful approach for studying these dynamic systems. The Martini model, which maps approximately two to four heavy atoms onto each coarse-grained bead, has been widely used to study membrane proteins [25, 26]. Compared with all-atom simulations, Martini is significantly more computationally efficient and can access longer time scales and larger systems [27]. The model also includes an extensive library of lipids that reproduce structural properties and phase behavior of lipid bilayers [28]. However, because Martini is an explicit-solvent model, simulations of transmembrane proteins with long IDRs can still be computationally expensive, as extended disordered regions require extensive sampling and large simulation boxes [15].

By representing the aqueous medium as a dielectric continuum, implicit-solvent models for IDRs and multi-domain proteins offer a considerable advantage in computational efficiency, enabling large-scale predictions of conformational ensembles and biomolecular condensates [29–32]. In recent years, several residue-level models have been developed to reproduce conformational properties and phase separation propensities of IDRs [33–40]. One such model is CALVADOS, which was optimized by tuning amino-acid-specific parameters against experimental data on the conformational properties of IDRs and multi-domain proteins, and captures propensities for biomolecular condensate formation [37, 41, 42]. While the model has been extended to nucleic acids [43] and polyethylene glycol [44], and applied to IDRs of membrane proteins [16], it has so far lacked an explicit representation of lipid membranes.

For lipids, implicit-solvent coarse-grained models were developed to capture generic structural and elastic properties of lipid bilayers at length and time scales that are difficult to access with explicit-solvent simulations [45–47]. The lipid model of Cooke et al. represents an archetypal phospholipid and has been used to study diverse membrane phenomena, including how lipid shape affects self-assembly [48, 49] and how proteins induce membrane budding [50]. More recent extensions of this class of models have incorporated additional features of biological membranes, including leaflet asymmetry and liquid-ordered/liquid-disordered phase coexistence [51, 52]. iSoLF increased the resolution of the Cooke model to five beads per lipid, making the lipid beads more comparable in size to those in one-bead-per-residue protein models, and was parameterized for POPC and DPPC bilayers [53]. In a later extension, iSoLF was combined with the residue-level protein model AICG2+ by tuning amino acid–lipid interactions against amino-acid transfer free energies, enabling membrane insertion and orientation to be captured with good agreement for several folded transmembrane proteins [54]. Finally, iSoLFv2 expanded the model to a larger set of lipid species by introducing electrostatic interactions and additional types of headgroup and tail beads, with parameters derived from all-atom simulations of lipid bilayers [55].

Here, we extend the CALVADOS framework to lipid membranes to study flexible transmembrane proteins and biomolecular condensates at the membrane–water interface. The resulting model, MEM-CALVADOS, captures nonspecific interactions between protein residues and lipid headgroups and tails, while retaining the sequence-dependent protein–protein interactions of CALVADOS. To reproduce membrane compositions commonly used in in vitro experiments, we introduce lipid models for POPC, DOPC, POPS, and DOPS, which differ in headgroup charge, tail length, and degree of unsaturation. Lipid parameters are refined using data from Martini and all-atom simulations, and the distance range of cross-interactions between hydrophobic residues and lipid tails is tuned to reproduce topology assignments and orientation predictions for a set of 21 transmembrane proteins [56, 57]. We validate the model against experimental transfer free energies of hydrophobic pentapeptides from the membrane to water and against an experimentally refined conformational ensemble of the human growth hormone receptor, a highly dynamic single-pass transmembrane protein with a long intracellular IDR [15, 58]. Finally, we apply the model to the signaling proteins LAT, Grb2, and Sos1, which assemble into membrane-bound clusters during T-cell activation, and examine how phosphorylation alters their interactions [8, 59, 60].

## Methods

### Protein Model

In MEM-CALVADOS, proteins are modeled as in CALVADOS 3 or AF-CALVADOS [32, 37, 41, 42]. Each amino acid is represented by a single bead, centered at the C_*α*_ position for IDRs and at the residue center of mass (COM) for folded domains of multi-domain proteins. In CALVADOS 3, folded domains are specified manually and COM–COM separations shorter than 0.9 nm are restrained at their initial values by harmonic potentials. AF-CALVADOS instead relies on structural models from the AlphaFold Database and uses predicted aligned errors (PAE) scores and predicted local distance difference (pLDDT) scores to identify and restrain folded domains through 12–10 Gō-type potentials [32]. Both approaches can be used in MEM-CALVADOS.

Bonded terms are modeled by a harmonic potential,

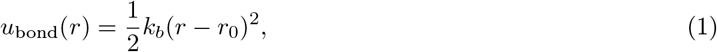

with *k*_*b*_ = 8033 kJ mol^−1^ nm^−2^. *r*_0_ is fixed at 0.38 nm in the case of both beads belonging to IDRs, while for folded domains it is determined by the distance between the COMs in the initial input structure.

Non-bonded interactions are modeled through the Ashbaugh-Hatch (AH) potential,

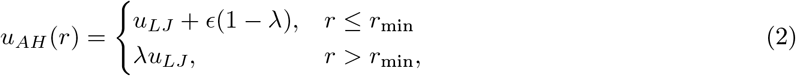

where *ϵ* = 0.8368 kJ mol^−1^ and *r*_min_ = 2^1*/*6^*σ. u*_LJ_ is the Lennard-Jones potential,

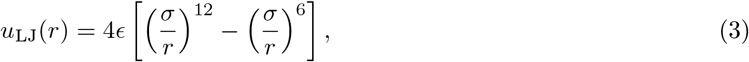

where *σ* = (*σ*_i_ + *σ*_j_)*/*2 and *λ* = (*λ*_i_ + *λ*_j_)*/*2 are the arithmetic averages of the respective parameters for beads

*i* and *j*. The AH potential is truncated and shifted at a cutoff of 2 nm.

The Debye-Hückel potential is applied to model salt-screened electrostatic interactions:

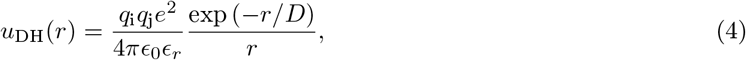

where *q* is the charge number, *e* is the elementary charge, 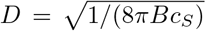 is the Debye length of an electrolyte solution of ionic strength *c*_*S*_, *B*(*ϵ*_*r*_) is the Bjerrum length, and *ϵ*_0_ is the vacuum permittivity.

*ϵ*_*r*_ is the dielectric constant of water modeled as dependent on the temperature, *T* , following the empirical relationship [61]:

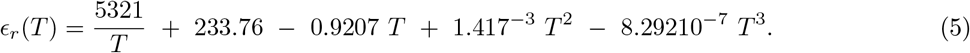

Fractional charges are assigned to histidine residues using the Henderson-Hasselbalch equation, assuming a p*K*_a_ of 6. *u*_DH_ is truncated and shifted at 4 nm. The potentials described so far define only interactions between amino acids; protein–membrane interactions are described separately below.

### Lipid Model

Building on iSoLFv2 [55], lipid models compatible with CALVADOS were developed in this work for 1-palmitoyl-2-oleoyl-sn-glycero-3-phosphocholine (POPC), 1,2-dioleoyl-sn-glycero-3-phosphocholine (DOPC), 1-palmitoyl-2-oleoyl-sn-glycero-3-phospho-L-serine (POPS), and 1,2-dioleoyl-sn-glycero-3-phospho-L-serine (DOPS). In iSoLFv2, these lipids are represented as linear chains of six beads with the first three beads describing the headgroup and the last three beads describing the hydrophobic tails (Figure 1). Consecutive beads are connected by harmonic bonds (Eq. 1), whereas chain stiffness is controlled by harmonic angle potentials along the lipid chain,

**Figure 1.**
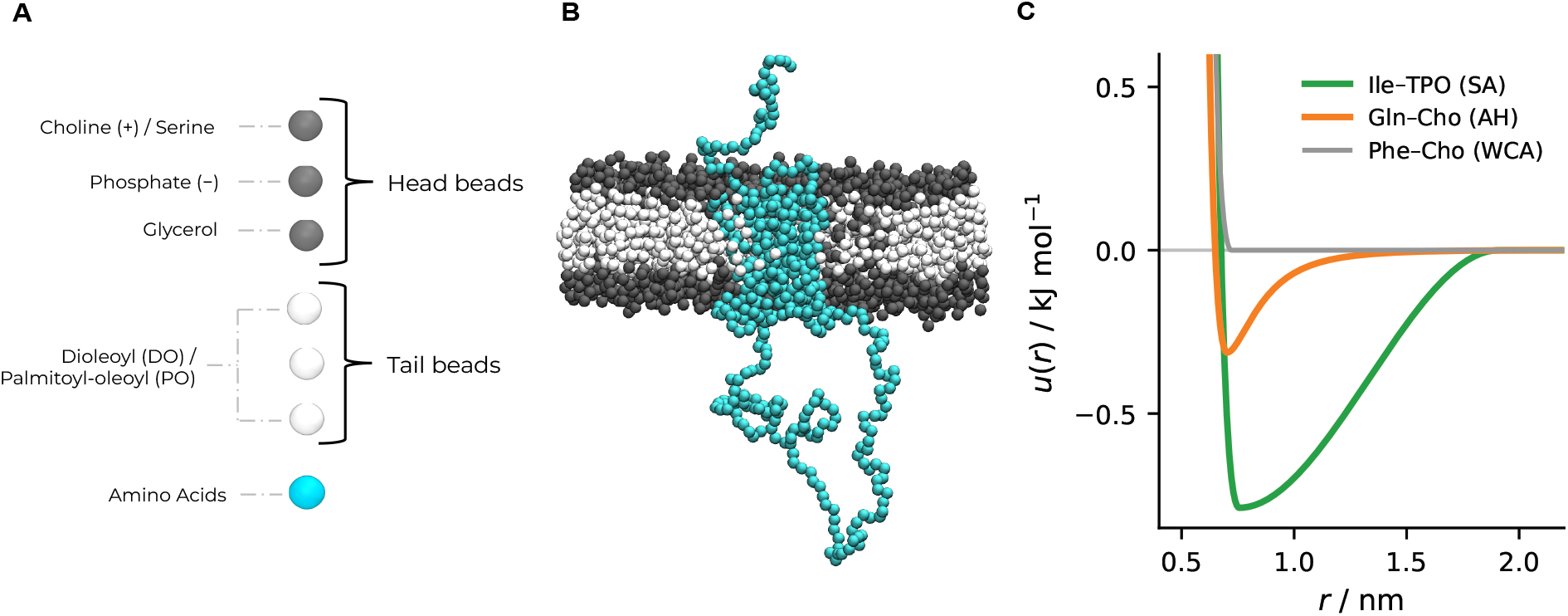
(A) Schematic representation of the different coarse–grained bead types of the MEM-CALVADOS model. (B) Snapshot of the transmembrane protein ADRA2A in a POPC bilayer. (C) Illustration of functional forms of the potentials used in the model: stretched attractive (SA), Ashbaugh-Hatch (AH), and Weeks-Chandler-Anderson (WCA). The examples shown are for Ile–TPO (green), Gln–Cho (orange), and Phe–Cho (gray). TPO is the palmitoyl–oleoyl tail bead.

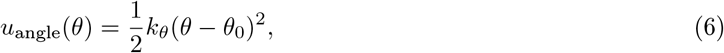

where *k*_*θ*_ is a force constant and *θ*_0_ is the equilibrium angle (Table S1). The terminal headgroup bead represents choline in PC lipids (Cho) or serine in PS lipids (Ser), whereas the two following beads are common to PC and PS lipids and represent phosphate (Pho) and glycerol (Mid). The hydrophobic region is represented by three tail beads, with distinct tail-bead types for palmitoyl-oleoyl and dioleoyl lipids. Tail–tail pairs interact through a stretched attractive (SA) potential which drives bilayer self-assembly,

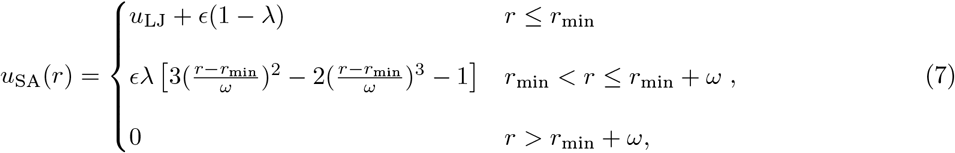

where *ω* = (*ω*_*i*_ +*ω*_*j*_)*/*2, *σ* = (*σ*_*i*_ +*σ*_*j*_)*/*2, and 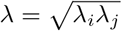 is the geometric average of the stickiness parameters. The SA potential decays smoothly to zero at *r*_min_+*ω* and its distance range is determined by the *ω* parameter. Tail–head pairs as well as all pair combinations involving Cho, Pho, and Mid interact through soft repulsions modeled with the Weeks–Chandler–Andersen (WCA) potential,

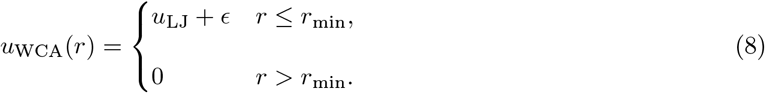

Interactions involving Ser beads are instead modeled using the AH potential truncated at 4 nm (Eq. 2). Cho and Pho beads are assigned unit positive and negative charges, respectively, and their mutual electrostatic interactions are modeled using Eq. 4.

Compared with the original iSoLFv2 models for POPC, DOPC, POPS, and DOPS, we introduced three modifications. First, force constants and equilibrium lengths of the two terminal tail-bond types were averaged, and the resulting common parameters were assigned to both bond types (Table S1). Second, the *ω* parameter was set to 1.2 nm for both palmitoyl-oleoyl and dioleoyl tail beads (Table S2). Third, the *λ* parameters of tail, Pho, and Cho beads were fine-tuned together with the *σ* parameters of the tail beads (Table S2). Starting values for lipid stickiness parameters were obtained from the iSoLFv2 *ϵ* values. For tail beads, the iSoLFv2 *ϵ* values were assigned directly as *λ*. For Cho, Pho, and Mid beads, the iSoLFv2 values were divided by four before assigning *λ*, to place headgroup stickiness values on the same scale as the CALVADOS stickiness parameters. Lipid Ser beads were assigned the same *λ* as the Ser amino-acid bead.

### Protein–Lipid Cross-Interactions

Nonbonded interactions between amino acid and lipid beads are described by the same potentials introduced for lipid–lipid interactions. The well depth of the potentials is scaled by the geometric mean of the stickiness parameters of the interacting amino acid and lipid beads while *σ* is obtained as the geometric mean. Amino acids were divided into three groups according to their expected affinity for lipid tails (Table S3 and Figure 1C)), using the transmembrane-tendency scale of Zhao and London, which quantifies the propensity of each residue to occur in *α*-helical transmembrane segments [62]. Amino acids with a transmembrane tendency less than or equal to that of Ala are treated analogously to Ser beads in PS lipids: they interact with headgroup beads through the AH potential (Eq. 2) and with tail beads through the WCA potential (Eq. 8). Amino acids with a transmembrane tendency greater than that of Ala are treated analogously to tail beads: they interact with headgroup beads through the WCA potential and with tail beads through the SA potential. Gly is treated as a special case. In CALVADOS, Gly has a large stickiness parameter [63], whereas its Wimley–White transfer free energy is similar to that of Ala. Because the large CALVADOS 3 stickiness of Gly reflects the compactness of Gly-rich IDR ensembles [64, 65] rather than an enhanced affinity for lipid bilayers, Gly–lipid interactions are modeled like Mid–lipid interactions, using WCA potentials with both headgroup and tail beads. The range of the SA cross-interactions is set by the arithmetic average of the *ω* value associated with the interacting tail and amino acid beads (Table S2).

### Molecular Simulations and Trajectory Analyses

Molecular dynamics simulations were performed using OpenMM v8.2 and the implementation of the model within the CALVADOS software package [66, 67]. Simulations were conducted using a Langevin integrator with a time step of 10 fs and a friction coefficient of 0.01 ps^−1^. Initial configurations were prepared by placing lipids in two leaflets on a regular grid in the *xy* plane, determining the number of lipids based on the initial box dimensions and the reference area per lipid reduced by 2%. The Monte Carlo membrane barostat implemented in OpenMM [68, 69] was used to allow isotropic fluctuations of the box dimensions in the membrane plane, with the external pressure and surface tension set to zero. The box length along *z* was fixed, and periodic boundary conditions were applied in all directions.

### Lipid Bilayers

Simulations of lipid bilayers used for parameter scanning and validation were run for 800 ns, saving frames every 1 ns and discarding the first 400 ns for equilibration. Simulations for the assessment of finite-size effects were run for 550 ns discarding the first 150 ns for equlibration. Simulations were performed at 297.15 K, unless stated otherwise. The initial dimensions of the bilayer patch were set to 25 nm ×25 nm, and to 12 nm ×12 nm to assess finite-size effects (Figure S1).

The area per lipid, APL, was calculated as

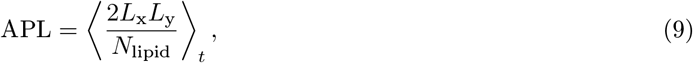

where *L*_x_ and *L*_y_ are box side lengths in the membrane plane, *N*_lipid_ is the total number of lipids, and

*⟨*… *⟩* _*t*_ denotes the time average. The phosphate–phosphate bilayer thickness, *d*_PP_, was computed as the time average of the instantaneous separation between the density peaks of the Pho beads in the two leaflets,

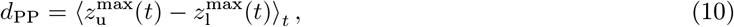

where 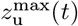 and 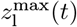 are the positions of the Pho density maxima in the upper and lower leaflet, respectively. Convergence of APL and *d*_PP_ was assessed by inspecting their cumulative averages (Figure S2). Standard errors were estimated through blocking analysis implemented in BLOCKING v0.1 (https://github.com/fpesceKU/BLOCKING).

### Single Transmembrane Proteins

For the simulation of single transmembrane proteins listed in Table S4, three 500-ns-long replicas were performed, preceded by 15-ns equilibration runs. Each protein was inserted in a POPC bilayer patch that spans the *xy*-plane of a box of initial dimensions 25 nm × 25 nm ×100 nm, with the exception of simulations of the human growth hormone receptor (GHR) and the epidermal growth factor receptor (EGFR), for which we used a box of initial dimensions 30 nm × 30 nm × 100 nm. AF-CALVADOS was used to simulate ADRA2A, RHO, TRPV4, OmpA, TOMM40, and VDAC1. All other proteins were simulated using harmonic restraints as in CALVADOS 3. For LAT, the restrained domain spanned residues 5–27. For GHR and EGFR, full-length structures were obtained from previous studies [5, 15], with CALVADOS 3 restraints applied to residues 52–287 and 639–863 for GHR, and residues 1–955 for EGFR. For protein structures obtained from the Protein Data Bank (PDB), harmonic restraints were applied over the entire sequence.

Membrane-inserted protein segments were identified from per-residue contacts between protein and lipid tail beads, calculated using the sigmoidal function

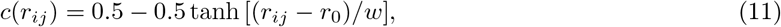

where *r*_*ij*_ is the distance between beads *i* and *j, r*_0_ = 1 nm, and *w* = 0.3 nm [31]. For each residue, contacts were summed over all lipid tail beads and averaged over the analyzed trajectory frames. Residue–tail contacts were averaged over the analyzed trajectory frames, smoothed using a five-residue moving average, and binarized using a threshold of one contact per frame. The resulting assignments of the transmembrane domains (TMDs) were compared with predictions from the machine-learning model DeepTMHMM v1.0, and the agreement was quantified using the intersection-over-union (IoU) metric,

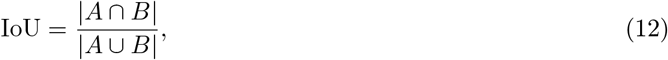

where *A* and *B* are the sets of residues assigned as transmembrane in the simulations and by DeepTMHMM, respectively.

Tilt angles of the TMDs were obtained using the segments assigned as transmembrane by DeepTMHMM. Tilt angles were computed as the angle between the *z*-axis, corresponding to the bilayer normal, and the principal axis of the TMD most closely aligned with the bilayer normal. Time-averaged tilt angles from simulations were compared with the top-ranked prediction from MemPrO [57], except for CD247, where the third-rank output was used because it was the highest-rank prediction in which the transmembrane helix spanned the membrane. The agreement between the mean tilt angle from simulations and the MemPrO reference was quantified by the absolute difference between the two values.

### WLXLL Peptides

Simulations of WLXLL peptides were performed starting from a bilayer patch of 12 nm × 12 nm, with a single peptide initially placed 2.6 nm away from the bilayer midplane. All 20 WLXLL peptides were simulated in two independent replicas at pH 7 and an ionic strength of 0.05 M. To model the N-acetylated peptides used in the experiments by Wimley and White [70], the N-terminal Trp bead was assigned a neutral charge, whereas the C-terminal Leu bead was assigned a negative charge to account for the carboxylate group. Each replica was run at both 303 and 323 K for at least 7.9 µs, and for over 50 µs for Met, Leu, Ile, and Phe at 323 K. The peptide–lipid potential energy was saved every 100 ps, and the first 100 ns were discarded for equilibration. The threshold separating membrane-bound and solution states was set to − 19.68 kJ mol^−1^, corresponding to the average position of the minima between the peaks in the peptide–lipid potential-energy distributions for the 20 amino acids. The bound fraction, *f*_bound_, was calculated as the fraction of frames for which the peptide–lipid potential energy was lower than the threshold. The free energy of partitioning from bulk solution to the bilayer was computed as

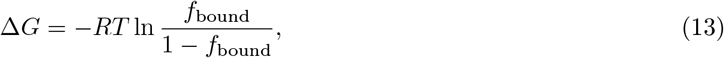

where *R* is the gas constant and *T* is the temperature. The mean Δ*G* and its standard error were estimated from the two independent replicas.

### Transmembrane Protein Assemblies

Systems comprising 16 copies of linker for activation of T-cells family member 1 (LAT; UniProt ID O43561) and 32 copies of both growth factor receptor-bound protein 2 (Grb2; UniProt ID P62993) and the proline-rich region of son of sevenless homolog 1 (Sos1; UniProt ID Q07889; residues 1117–1319) were simulated in a cuboidal box with initial dimensions 30 nm × 30 nm × 150 nm. The initial structure for LAT was extracted from a MEM-CALVADOS simulation of a single copy of the protein in a lipid bilayer, where the intracellar IDR extended away from the membrane. For LAT, residues 5–27 were restrained as in CALVADOS 3 and residue 15 was initially positioned at the midplane of a bilayer of 2,688 DOPC molecules spanning the *xy*-plane. Grb2 was simulated using AF-CALVADOS, with PAE and pLDDT data obtained from the AlphaFold Database entry AF-P62993-F1.

Phosphorylated LAT (pLAT) was modeled by replacing Tyr residues 131, 170, 190, and 225 with phosphorylated tyrosine (pTyr). The pTyr bead was assigned *σ* = 0.703 nm, derived by assuming a volume increase of 0.041 nm^3^ relative to Tyr [44, 71], and a charge of −1.97, calculated at pH 7.4 from the Henderson–Hasselbalch equation using *pK*_a_ = 5.83 [72]. Its *λ* parameter was set to 0.581 by applying the shift Δ*λ* = −0.37 to the CALVADOS 3 value for Tyr, following the shift used by Rauh et al. for pSer and pThr [44]. In contrast to Tyr, which interacts with lipid tails through the SA potential and with head-group beads through WCA interactions, pTyr interacts with lipid tails through WCA interactions and with headgroup beads through the AH potential.

Simulations were performed in five independent replicas of 1.2 µs each at pH 7.4, 293.15 K, and an ionic strength of 0.15 M. Frames were saved every 0.5 ns and the first 300 ns were discarded as equilibration based on the time series of LAT–Sos1 contacts. Concentration profiles were calculated as a function of the *z*-component of the protein COM after centering each trajectory frame so that the bilayer midplane is located at *z* = 0 nm. Per-residue protein–protein contacts were calculated using Eq. 11 and averaged over all protein pairs and analyzed trajectory frames.

## Results and Discussion

### Fine-tuning and Validation of Lipid Model

After modifying the iSoLFv2 bonded and nonbonded interactions as described in Methods, we fine-tuned selected lipid parameters using as reference data APL and *d*_PP_ values from Martini and all-atom simulations [28, 75–77]. For optimization of the tail-bead parameters, we used simulations of single-component DOPC and POPC bilayers and compared the resulting membrane properties with Martini 2 and Martini 3 reference data (Figure S3) [28]. For Pho and Cho, we used PS-containing bilayers because only their AH interactions with Ser beads depend on *λ*. The *λ* value for Pho was optimized using single-component DOPS and POPS bilayers with Martini reference data (Figure S4) [28], and the *λ* value for Cho using two-component POPS/POPC and DOPS/DOPC bilayers with all-atom reference data (Figure S5 and Figure 2D) [75–77]. The optimization was performed stepwise. We first optimized the *σ* and *λ* parameters of the tail beads by grid search. Using the optimized tail-bead parameters, we then scanned the *λ* parameter of Pho beads, followed by the *λ* parameter of Cho beads. We found that changes of less than 3% relative to the initial parameters were sufficient to improve the agreement with the reference data (Figures S3–S5). For tail beads, the optimized parameters suggest that dioleoyl beads are better described by a smaller *λ* and a larger *σ* than palmitoyl–oleoyl beads (Figure S3), consistent with the looser packing expected for more unsaturated lipid tails.

**Figure 2.**
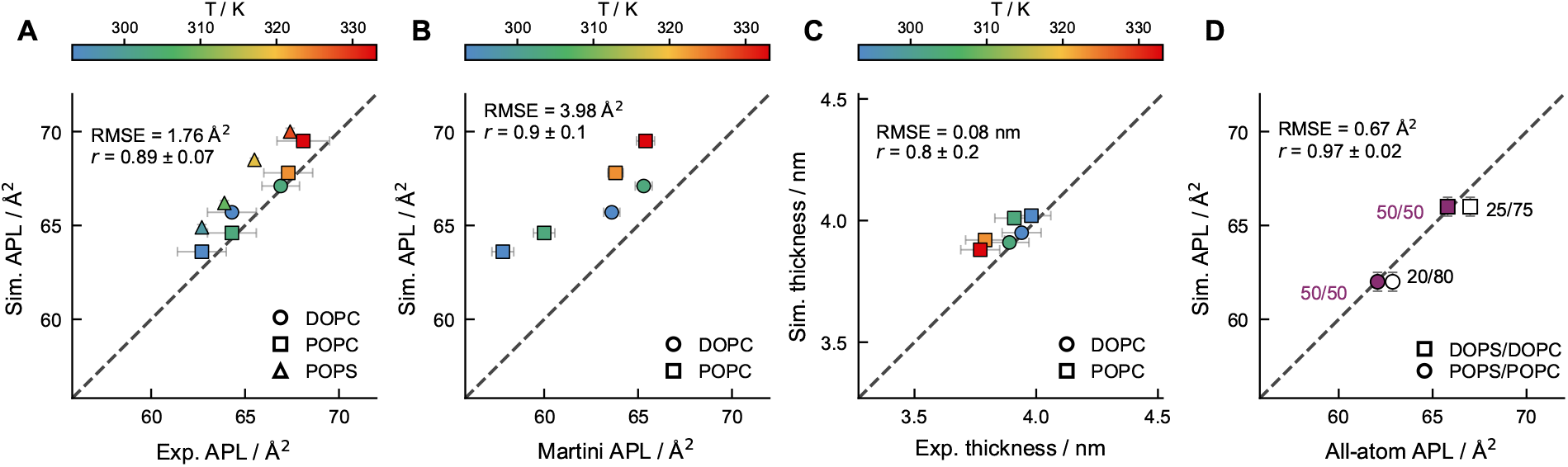
Validation of the lipid model. (A, B) Comparison of area per lipid (APL) from our model with(A)experimental data and (B) Martini 3 simulations at different temperatures for single-component POPC (293.15, 303.15, 323.15, and 333.15 K), DOPC (293.15 and 303.15 K), and POPS (298.15, 308.15, 318.15, and 328.15 K) bilayers [28, 73, 74]. (C) Comparison of phosphate–phosphate bilayer thickness from our model and Martini 3 simulations at different temperatures for POPC (293.15, 303.15, 323.15, and 333.15 K) and DOPC (293.15 and 303.15 K) bilayers [28]. (D) Comparison of APL from our model and all-atom simulations for mixed DOPS–DOPC and POPS–POPC bilayers at 310 and 298 K, respectively, and at different molar ratios [75–77]. *r* is the Pearson correlation coefficient, with uncertainty estimated as the standard deviation over 10^4^ bootstrap samples, and RMSE is the root-mean-square error.

To validate the optimized lipid model, we simulated PC and PS lipids at different temperatures (Figure 2). Comparison with reference data from experiments [28, 73, 74] and Martini 3 simulations [28] shows that the model recapitulates the temperature dependence of APL and *d*_PP_ (Figure 2A,C). Notably, the model captures the APL of PC lipids more accurately than Martini 3 (Figure 2B), which was previously shown to underestimate this property by approximately 0.03 nm^2^ [28].

### Calibration of Protein–Lipid Interaction Parameter

In the model, self-assembly of lipid bilayers is driven by attractive interactions between tail beads. We used the same SA potential (Eq. 7 and Figure 1C) to describe interactions between lipid tail beads and amino acids with greater transmembrane tendency than Ala on the Zhao–London scale [62]. In the SA potential, *ω* controls the range of the attraction and is therefore expected to influence both membrane insertion and the orientation of TMDs. We thus set out to tune this parameter for protein–lipid cross-interactions using simulations of transmembrane proteins. To this end, we collected a set of 21 membrane proteins with diverse topologies, including single-pass *α*-helical proteins, multipass *α*-helical proteins, *β*-barrel proteins, and multi-subunit complexes (Table S4). The input structures comprise nine AlphaFold Database models, two previously published full-length models of single-pass receptors [5, 15], and ten PDB structures previously used to benchmark the MemPrO tool [57]. For each protein or complex, we performed three independent 500-ns simulations in a POPC bilayer using *ω* values of 1.0, 1.1, and 1.2 nm, corresponding to increasing distance ranges for the SA interaction between amino acids with greater transmembrane tendency than Ala and lipid tail beads. For each value of *ω*, we quantified the agreement with predicted membrane insertion and orientation (Figure 3). Membrane-inserted regions were identified from residue–tail contacts in the simulations (Figure S6A–C) and compared with DeepTMHMM transmembrane-segment assignments using the IoU metric (Figure 3A,C). TMD tilt angles were averaged over the simulation trajectories (Figure S6D– F) and compared with orientation predictions from MemPrO using the absolute error (Figure 3B,D) [57]. DeepTMHMM is a deep-learning sequence-based method for residue-level topology prediction of both *α*-helical and *β*-barrel transmembrane proteins [56], whereas MemPrO is a method that uses a Martini-based mean-field scoring function to predict membrane insertion depth and orientation of proteins from their structures [57].

**Figure 3.**
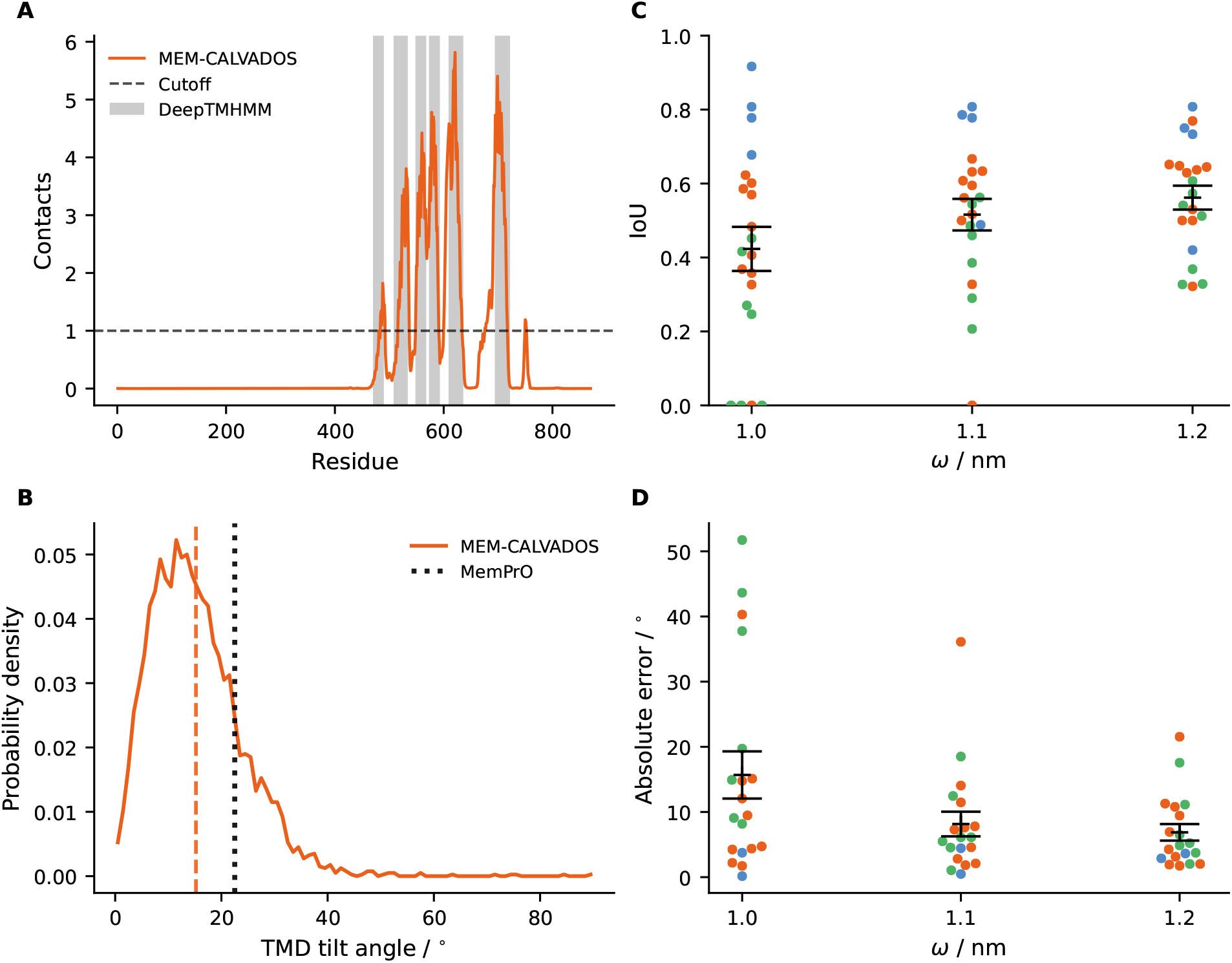
Calibration of the *ω* parameter for hydrophobic amino acids. (A) Per-residue contacts between TRPV4 and lipid tail beads from simulations in a DOPC bilayer with *ω* = 1.1 nm (orange). Gray shaded regions indicate transmembrane segments predicted by DeepTMHMM, and the dashed black line indicates the threshold of one contact per frame used to assign membrane-inserted residues from the contact profile.(B) Distribution of TRPV4 TMD tilt angles from simulations with *ω* = 1.1 nm (orange). The orange vertical line indicates the simulation average, and the black dotted line indicates the MemPrO prediction. (C) Agreement between transmembrane-segment assignments from MEM-CALVADOS simulations and DeepTMHMM, quantified by the IoU for different *ω* values. (D) Absolute error between simulated TMD tilt angles and MemPrO orientation predictions for different *ω* values. Each circle in panels C and D represents one protein, with the color indicating the TMD architecture: single-pass *α*-helical (blue), multipass *α*-helical (orange), and *β*-barrel (green). Black markers and error bars indicate the mean and standard error of the absolute errors across proteins at each *ω* value.

For single-pass *α*-helical proteins, the model is in good agreement with both topology and orientation predictions. In GHR, EGFR, LAT, and CD247, the membrane-inserted regions obtained from simulations closely match the transmembrane segments predicted by DeepTMHMM, and the TMD tilt angles agree well with MemPrO predictions for all tested *ω* values (Figures S7–S8 and S12–S13). The long disordered intracellular regions present in these proteins do not partition into the membrane. However, the N-terminal signal peptide in the simulated full-length GHR structure forms contacts with the membrane for *ω >* 1.0 nm.

For *ω* = 1.1 nm, the average number of contacts between signal peptide and tail beads is approximately fourfold lower than that of the transmembrane helix, whereas at *ω* = 1.2 nm the signal peptide is stably associated with the membrane, with contacts comparable to those of the transmembrane helix (Figure S7A). Although this signal peptide is cleaved before the mature receptor reaches the plasma membrane, it was included in the construct studied by Kassem et al. and did not form long-lived contacts with the membrane in their Martini simulations [15]. Its stable membrane association at *ω* = 1.2 nm therefore suggests that this interaction range results in overly attractive residue–tail interactions.

Multipass *α*-helical proteins are also generally well captured. MEM-CALVADOS results for the seven-transmembrane-helix bundles of ADRA2A and rhodopsin agree with the DeepTMHMM topology assignments and MemPrO orientation predictions for *ω >* 1.0 nm, whereas ADRA2A detaches from the membrane at *ω* = 1.0 nm (Figures S8 and S13–S14). Simulations of the full-length TRPV4 monomer, which has a different transmembrane architecture comprising six helices rather than a GPCR-like bundle, also capture membrane insertion and orientation in agreement with the reference predictions (Figures 3A and Figures S8E and S14B). Similarly, the multipass *α*-helical TMDs of 5CFB, 5VPN, 5XU1, 8TOL, and 4IKV remain inserted throughout the simulations and show good agreement with the predicted orientations (Figures S10–S11 and S16–S17). By contrast, simulations of smaller isolated two-helix systems are in weaker agreement with the reference data. The *α*-helical transmembrane dimer of the p75 neurotrophin receptor (PDB 2MJO) partitions into the membrane only at *ω* = 1.2 nm (Figures S10 and S16) whereas the helix hairpin of F_1_F_o_ ATP synthase subunit c (PDB 1A91) remains inserted for *ω >* 1.0 nm but its mean tilt angle is overestimated relative to the reference (Figures S10 and S16). We note, however, that at *ω* = 1.1 nm PDB 1A91 also samples a less populated state with a more perpendicular orientation relative to the membrane (Figures S6E and S16B). These discrepancies may reflect limitations in the residue-level resolution of the model, as isolated transmembrane helices can depend on specific side-chain–lipid interactions, including those involving aromatic residues, for stable insertion and orientation [78].

The *β*-barrel proteins are particularly informative for identifying the lower bound of the hydrophobic residue–tail interaction range. Although all proteins are initially inserted in the membrane, most *β*-barrels detach from the bilayer during simulations with *ω* = 1.0 nm. For *ω >* 1.0 nm, all *β*-barrel proteins remain inserted in the membrane (Figures S9 and S11). For most of these proteins, the simulated orientations are close to the MemPrO predictions with absolute error ≤6^*◦*^, although larger deviations are observed for BamA from both *E. coli* and *N. gonorrhoeae* (Figures S14–S15 and S18). The larger trimeric *β*-barrel complex TolC (PDB 1EK9), which contains more than 1,000 residues and forms a single membrane-spanning barrel from three monomers, also remains membrane-associated for *ω >* 1.0 nm, with an absolute error of 4.5^*◦*^ in the average TMD tilt angle (Figures S10 and S17). Thus, *ω* = 1.0 nm is insufficient to stabilize *β*-barrel insertion, whereas larger values result in stable membrane association for most *β*-barrel systems. Taken together, these results identify *ω* = 1.1 nm as the shortest interaction range that stabilizes membrane insertion of multipass *α*-helical proteins and *β*-barrels. Given the comparable mean IoU and mean absolute error in tilt angle at *ω* = 1.1 and 1.2 nm, we select *ω* = 1.1 nm to minimize spurious membrane association, such as that observed for the signal peptide of GHR at *ω* = 1.2 nm.

### Validation Against Wimley–White Transfer Free Energies

In combining iSoLFv2 and CALVADOS, we assumed that hydrophobic residue–tail interactions can be described by the same SA potential used for tail–tail interactions, whereas polar residue–headgroup interactions are described by the AH potential. The strengths of these interactions are scaled by combining amino-acid and lipid-bead stickiness parameters from CALVADOS and iSoLFv2 through geometric averages. The residues to which we assign attractive interactions with lipid tails are V, M, I, L, F, Y, and W. This selection is based on their propensity to occur in *α*-helical transmembrane segments, as quantified by the transmembrane-tendency scale [62]. To validate this approach, we compared simulated transfer free energies from bilayer to water, Δ*G*, for WLXLL pentapeptides, where X denotes each of the 20 amino acids, with the experimental values reported by Wimley and White [70].

We estimated the Δ*G* values from unbiased simulations using the relative populations of the membrane-associated and solution states, identified from their peptide–lipid interaction energies (Figure 4A and Eq. 13). To increase the rate of exchange between these states and improve sampling, we performed simulations at 303 and 323 K. The Δ*G* values at 303 K show a small systematic increase relative to those at 323 K, with an RMSE of 1.5 kJ mol^−1^, while the two sets of values are strongly correlated (*r* = 0.99 0.09; Figure 4B), consistent with the weak temperature dependence reported for hydrophobic peptides in a previous all-atom study [80]. At 303 K, converged Δ*G* values were obtained for peptides with X = E, D, K, R, Q, P, N, H, T, A, S, V, G, C, and L; at 323 K, convergence was also achieved for peptides with X = M and I.

**Figure 4.**
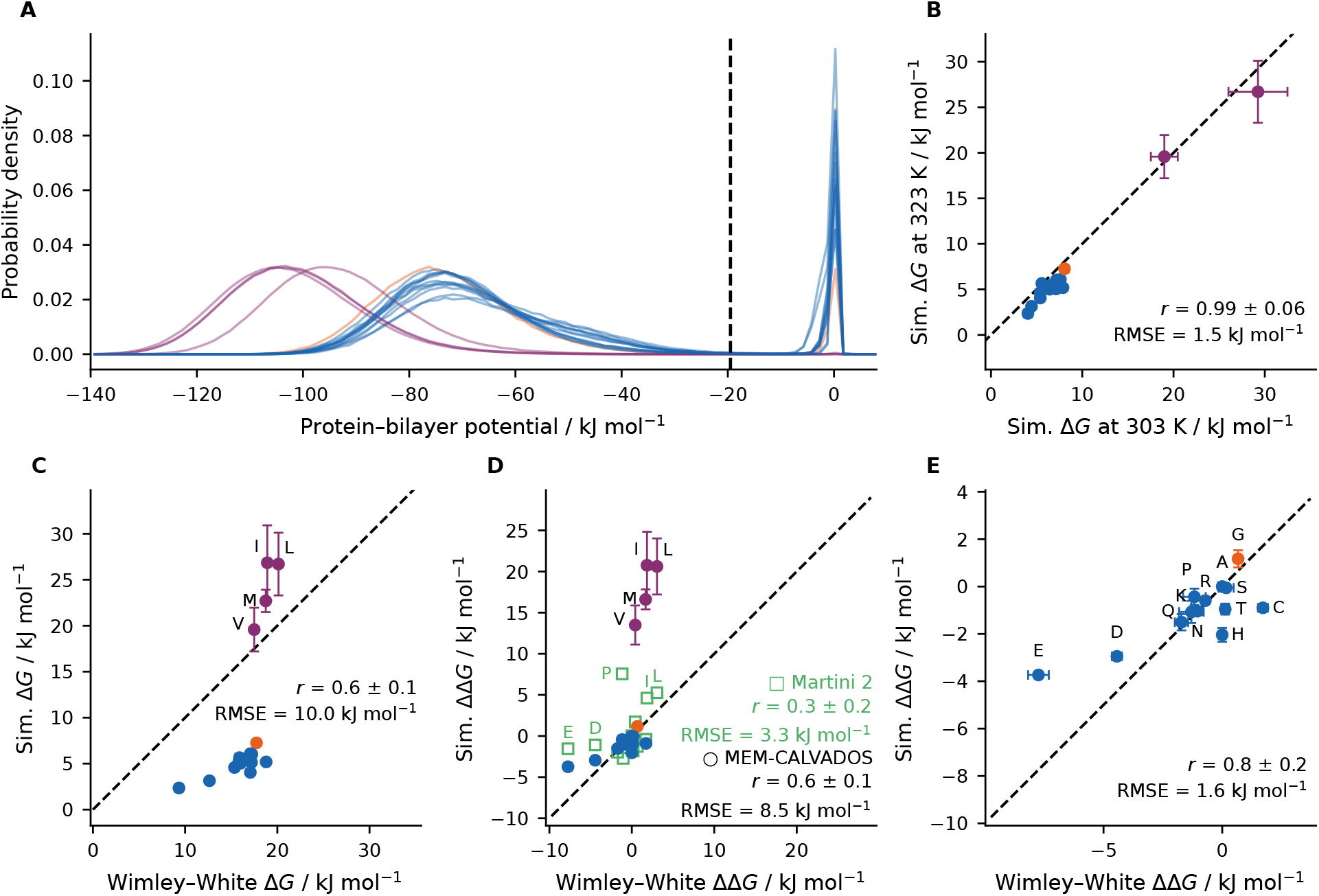
(A) Probability density distributions of the total peptide–tail interaction energy for the 20 WLXLL peptides. The black dotted line indicates the average position of the minimum separating the bound and unbound peaks, which was used as the threshold to define bound and unbound states. (B) Correlation between transfer free energies, Δ*G*, from MEM-CALVADOS simulations performed at 303 and 323 K (Eq. 13). (C) Correlation between Δ*G* from simulations at 323 K and the corresponding experimental values [70].(D) Correlation of relative transfer free energies, ΔΔ*G* = Δ*G*_WLXLL_ − Δ*G*_WLALL_, from MEM-CALVADOS simulations at 323 K or Martini 2 simulations at 300 K [79] and the corresponding experimental values [70], for the subset of peptides available for both models. Martini 2 values are shown in green.(E) Correlation of ΔΔ*G* values from MEM-CALVADOS simulations at 323 K and experimental values [70] for all peptides in which the central amino acid interacts with lipid tails through WCA interactions. Colors indicate peptides with Gly (orange), amino acids interacting with tails through SA potential (purple), and amino acids interacting with tails with WCA potential (blue). MEM-CALVADOS data in B–E are mean *±* standard error of the mean estimated from two independent replicas. *r* is the Pearson correlation coefficient, with uncertainty estimated as the standard deviation over 10^4^ bootstrap samples, and RMSE denotes the root-mean-square error.

MEM-CALVADOS reproduces the overall ranking of the Wimley–White transfer free energies. Across all peptides, the correlation between simulated and experimental Δ*G* values is *r* = 0.6 *±* 0.1, and the transfer free energies of peptides with X = V, M, I, and L are overestimated with an RMSE of 5.6 kJ mol^−1^ (Figure 4C). In contrast, the model systematically underestimates Δ*G* for peptides containing residues that are not assigned attractive interactions with lipid tails, with an RMSE of 11.0 kJ mol^−1^. This pronounced energetic separation between hydrophobic and polar residues likely reflects the binary treatment of residue–tail interactions in the model, which are either attractive or purely repulsive based on the ranking of the amino acid in the transmembrane-tendency scale [62]. A smoother dependence of the interaction strength on the propensity to occur in TMDs may reduce this separation and could be considered in future refinements of the model.

In a previous study, Martini 2 transfer free-energy differences relative to WLALL, ΔΔ*G* = Δ*G*_WLXLL_ −Δ*G*_WLALL_, were calculated using alchemical thermodynamic cycles [79]. In that work, peptides with X = R and K were simulated with a protonated C terminus to mimic the Wimley–White experiments at pH 2, whereas we used a negatively charged C terminus for all peptides. We therefore excluded R and K and compared the remaining peptides common to both studies with ΔΔ*G* derived from the Δ*G* values reported by Wimley and White for pH 8. For this subset, MEM-CALVADOS reproduces the experimental ΔΔ*G* trends with *r* = 0.6*±* 0.1 and an RMSE of 8.5 kJ mol^−1^, whereas Martini 2 gives a lower correlation, *r* = 0.3*±* 0.2, but a smaller RMSE of 3.3 kJ mol^−1^ (Figure 4D). When the comparison is restricted to peptides with X having a transmembrane tendency no greater than that of Ala, MEM-CALVADOS is in excellent agreement with the Wimley–White ΔΔ*G* values, with *r* = 0.8 *±*0.2 and an RMSE of 1.6 kJ mol^−1^ (Figure 4E).

### Conformational Properties of the Growth Hormone Receptor

Having validated the residue–lipid cross-interactions against experimental transfer free energies, we next tested the ability of MEM-CALVADOS to reproduce the conformational properties of the full-length human growth hormone receptor (GHR). GHR is a single-pass transmembrane receptor of the class 1 cytokine receptor family and comprises a folded extracellular domain (ECD), an *α*-helical TMD, and a long intrinsically disordered intracellular domain (ICD). Its conformational ensemble was previously characterized using an integrative approach that combined SAXS and SANS measurements of the receptor in POPC nanodiscs with NMR spectroscopy, X-ray diffraction, and Martini simulations in a POPC bilayer [15]. In the Martini simulations, protein–water interactions were strengthened by 10% to counteract the tendency of the force field to overestimate IDR compaction [81]. The ensemble generated by unbiased molecular dynamics was subsequently reweighted against the experimental SAXS profile. The resulting experimentally refined ensemble provides distributions of the radii of gyration of the full receptor and its individual domains, as well as the orientations of the transmembrane helix and extracellular domain relative to the membrane normal. For direct comparison, we simulated the same full-length GHR construct under the conditions of the Martini simulations, i.e., 310 K and an ionic strength of 0.15 M. The construct includes the N-terminal signal peptide and a green fluorescent protein tag attached to the C-terminus to facilitate purification [15] (Figure 5A). The *R*_*g*_ distribution of the full-length construct is in excellent agreement with the experimentally refined Martini ensemble, with a relative error of −0.01% in the mean *R*_*g*_. By comparison, the Martini ensemble before reweighting overestimated the mean *R*_*g*_ by 10%. MEM-CALVADOS also accurately reproduces the compaction of the ICD, both with and without GFP, while it underestimates the *R*_*g*_ of the ECD, including the signal peptide, by 4% (Figure 5B).

**Figure 5.**
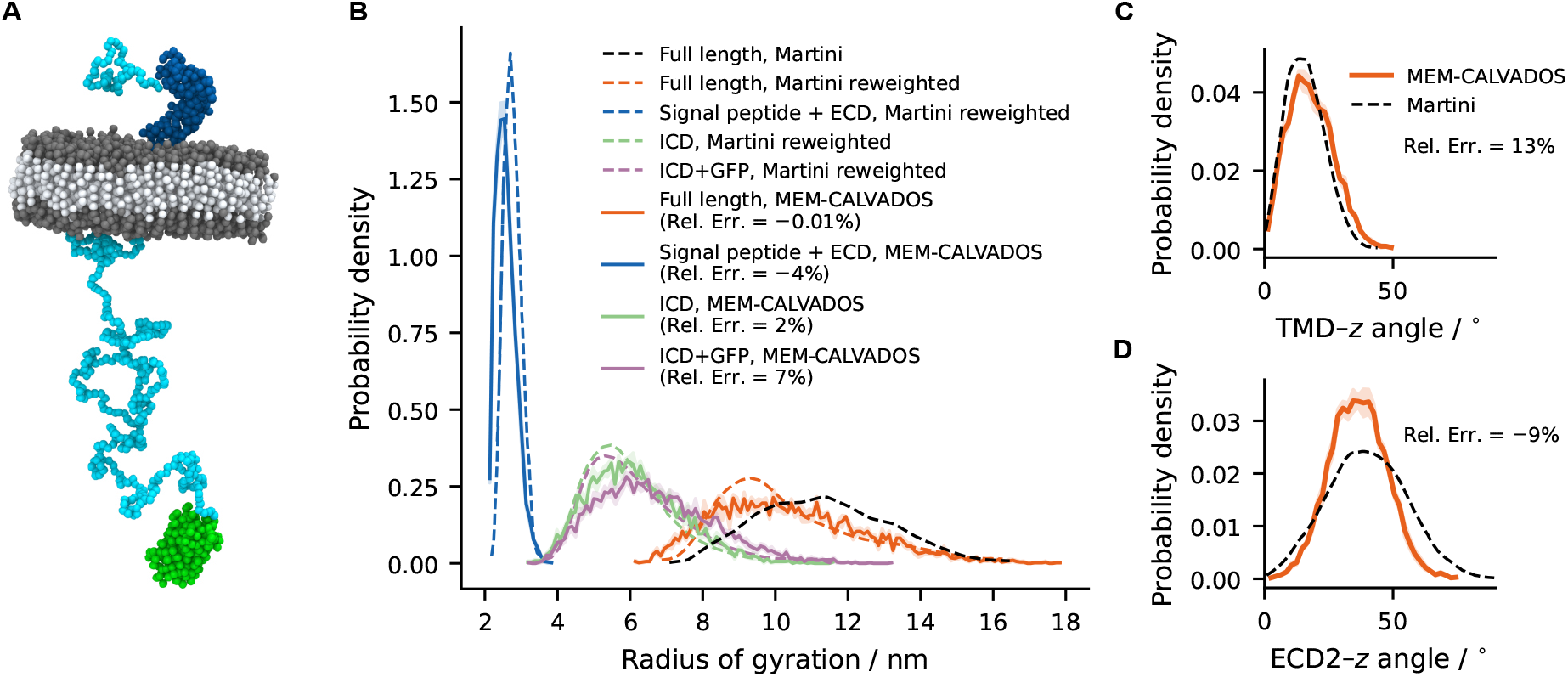
Conformational properties of GHR. (A) Snapshot of the full-length construct of GHR studied by Kassem et al. [15] including the intrinsically disordered extracellular signal peptide and intracellular domain (cyan), the folded extracellular domains D1 and D2 (blue), and the green fluorescence protein attached to the C-terminus (green). The protein is embedded in a bilayer of 2,856 POPC lipids, shown partially for clarity.(B) Distributions of the radius of gyration, *R*_*g*_, from MEM-CALVADOS simulations (solid lines) compared with Martini simulations from Kassem et al. [15]. Dashed colored lines show Martini distributions reweighted against SAXS data, and the black dashed line shows the Martini distribution for the full-length construct before reweighting. (C) Distribution of the TMD tilt angle from MEM-CALVADOS simulations (solid line) and SAXS-reweighted Martini simulations from Kassem et al. (dashed line). (D) Distribution of the angle between the principal axis of the extracellular D2 domain and the bilayer normal from MEM-CALVADOS simulations (solid line) and SAXS-reweighted Martini simulations (dashed line). For the MEM-CALVADOS data, solid lines show the mean over five replicas, and shaded areas indicate the standard error of the mean. Relative errors are reported for the mean *R*_*g*_ of MEM-CALVADOS with respect to the reweighted Martini ensemble.

The TMD tilt-angle distribution (Figure 5C) and the orientation of the D2 principal axis relative to the bilayer normal (Figure 5D) also agree well with the reweighted Martini ensemble, with relative errors in the mean angles of 13% and− 9%, respectively. These results indicate that MEM-CALVADOS captures both the predominantly perpendicular orientation of the TMD, consistent with X-ray diffraction data, and the orientational flexibility of the ECD–TMD linker observed in Martini simulations [15].

To assess whether the similarities between MEM-CALVADOS and Martini observed for GHR also extend to protein–membrane interactions, we calculated contacts between protein residues and both lipid tails and phosphate beads, and compared them with Martini trajectories before reweighting (Figure S19). The two models show remarkably similar contact profiles for the transmembrane region and residues in the extracellular linker. The main discrepancy concerns the Box1 motif, which forms persistent membrane contacts in Martini but not in MEM-CALVADOS (Figure S19).

These results show that MEM-CALVADOS reproduces the conformational properties of full-length GHR without requiring reweighting against experimental data. The agreement obtained for the ICD is consistent with previous CALVADOS benchmarks for intrinsically disordered proteins [41], while our simulations further show that the model captures GHR–membrane interactions, as reflected in receptor orientations and residue-resolved lipid contacts. Moreover, these observables are obtained at a low computational cost, with a single 515-ns simulation requiring approximately 14 h on a single GH2000 GPU using CUDA and providing stable estimates of the reported quantities (Figure S20).

### Clustering of T-cell Signaling Proteins

To assess whether our model reproduces the clustering of signaling proteins at the bilayer–water interface, we simulated a system comprising LAT, Grb2, and the proline-rich region of Sos1. LAT is a transmembrane adaptor protein consisting of a short extracellular domain, a single transmembrane helix, and an intrinsically disordered intracellular domain. During T-cell activation, LAT is phosphorylated by the tyrosine kinase ZAP70, and its phosphotyrosines are key to the recruitment of downstream signaling proteins. One of its principal binding partners is Grb2, a folded adaptor protein with an SH3–SH2–SH3 domain architecture. The SH2 domain binds distal phosphotyrosines in LAT, whereas the two SH3 domains bind proline-rich motifs in Sos1, thereby forming a multivalent interaction network promoting protein clustering [8, 59, 60].

Su et al. reconstituted the system using the cytoplasmic portion of LAT, fluorescently labeled and tethered to a supported lipid bilayer with a surface density of 300 molecules µm^−2^. Using total internal reflection fluorescence microscopy, micrometer-sized clusters of phosphorylated LAT (pLAT) formed in the presence of Grb2 and the proline-rich region of Sos1. Unphosphorylated LAT did not cluster, and substitution of three distal Tyr residues with Phe suppressed clustering [60].

To model this system, we simulated 16 copies of full-length LAT, with their transmembrane helices embedded in a DOPC bilayer, together with 32 copies of full-length Grb2 and the proline-rich region of Sos1. These conditions correspond to a LAT surface density of 17 × 10^3^ molecules µm^−2^ and a solution concentration of 45 µM for Grb2 and Sos1. In our simulations, Grb2 and Sos1 freely diffuse in the compartments above and below the bilayer, which are connected through periodic boundary conditions. LAT is instead embedded in the membrane with its intracellular domain positioned in the lower half of the simulation box (Figure 6A,B). Protein concentration profiles along the bilayer normal show that Sos1 preferentially localizes to the bilayer–water interface, with a concentration peak in the region occupied by the LAT intracellular domains (Figure 6D). Phosphorylation of LAT further enhances this enrichment, as quantified by the excess number of Sos1 chains, Δ*N* , in the intracellular region, −20 nm *< z*_COM_ *<* 0 nm, relative to the corresponding extra-cellular region, 0 nm *< z*_COM_ *<* 20 nm. Δ*N* increases from 1.0 *±* 0.2 for the system with unphosphorylated LAT to 3.5 *±* 0.3 in the presence of pLAT (Figure S21A). In contrast, Grb2 remains approximately equally distributed between the two compartments in both systems (Figure 6C,D).

**Figure 6.**
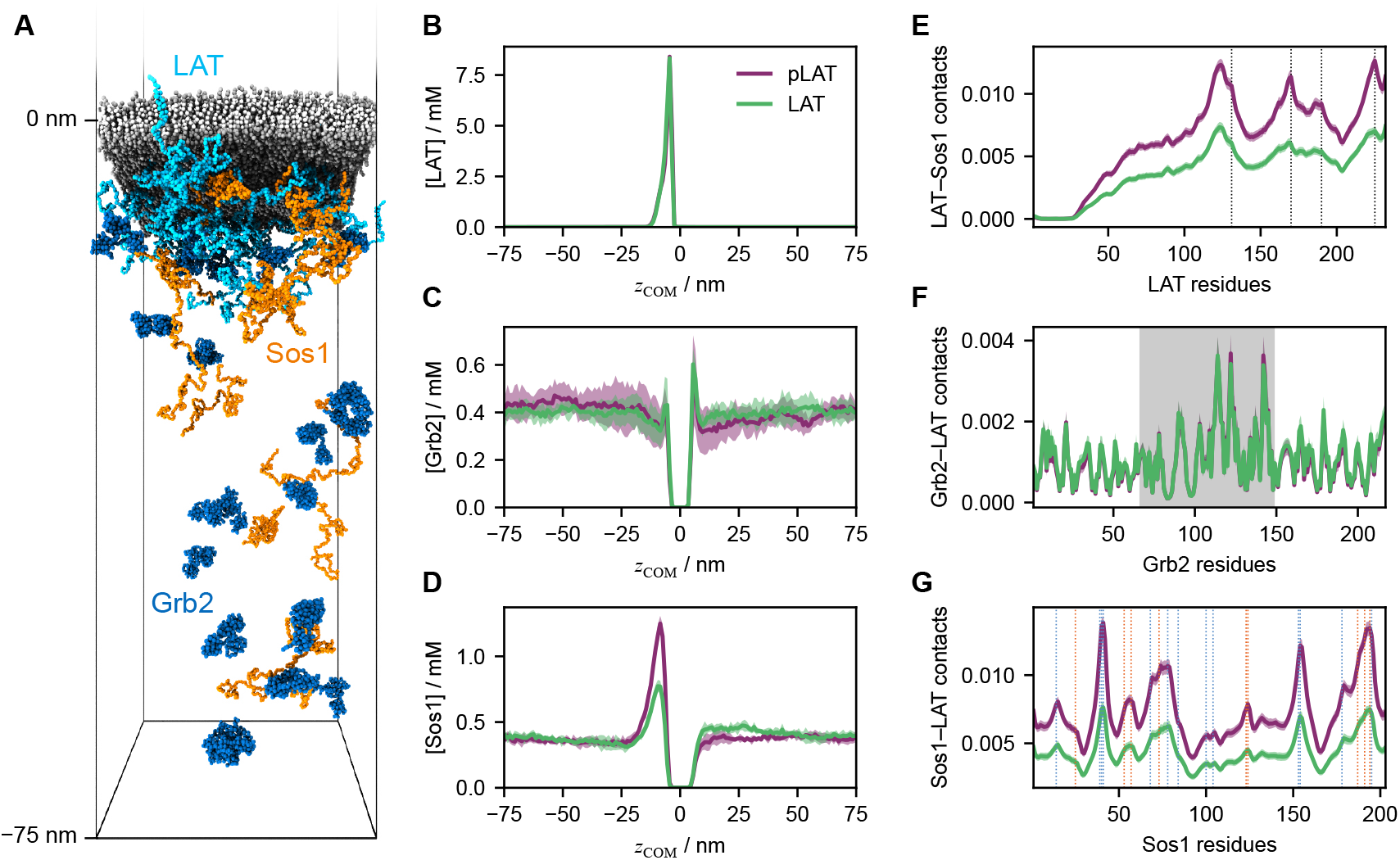
Clustering of LAT, Grb2, and Sos1. (A) Snapshot of the simulated system comprising a lipid bilayer of 2,688 DOPC molecules, 16 copies of the transmembrane protein LAT (cyan), and 32 copies of both cytosolic proteins Grb2 (blue) and Sos1 (orange) in a cuboidal box with a fixed length of 150 nm along *z*. The snapshot shows the lower half of the box where the intracellular domain of LAT is positioned. (B–D) Concentration profiles of (B) LAT, (C) Grb2, and (D) Sos1 in the systems with phosphorylated (purple) and unphosphorylated (green) LAT as a function of the *z*-component of the center of mass, *z*_COM_, of the corresponding protein. (E–G) Per-residue profiles of (E) LAT contacts with Sos1, (F) Grb2 contacts with LAT, and (G) Sos1 contacts with LAT in the systems with phosphorylated (purple) and unphosphorylated (green) LAT. Dotted lines in panel E indicate the positions of the phosphorylated tyrosines. The gray shaded area in panel F indicates the SH2 domain. Dotted lines in panel G indicate the positions of arginine (blue) and lysine (orange) residues in Sos1. Solid lines are means over five independent replicas, and green and purple shaded areas indicate the standard error of the mean.

The increased accumulation of Sos1 on the intracellular side is reflected in a 66 *±* 7% increase in LAT– Sos1 residue–residue contacts upon LAT phosphorylation (Figure S21B). The corresponding contact profiles are characterized by interactions between regions surrounding the distal tyrosines of LAT and Arg/Lys-rich regions of Sos1, which are observed with unphosphorylated LAT and are strongly enhanced in the phosphorylated system (Figure 6E,G). Conversely, Grb2–LAT and Grb2–Sos1 contacts are negligibly affected by LAT phosphorylation (Figure 6F and Figure S21C), indicating that the model does not fully capture the role of Grb2 in cluster formation. The Grb2–LAT contact profile is nevertheless consistent with the expected binding mode, with contacts predominantly involving the SH2 domain of Grb2, which has been shown to recognize pY-X-N-X motifs present in LAT [59, 82].

We note that, because of the high LAT surface density in our simulations, we do not expect to observe a further enhancement in clustering upon phosphorylation. Reproducing the experimental LAT surface density with 16 LAT molecules would require bilayer patches more than an order of magnitude larger in each lateral dimension, which would substantially limit the accessible simulation time. Instead, our simulations probe how phosphorylation modulates the underlying interaction network at the bilayer–water interface. Our results indicate that phosphorylation primarily enhances LAT–Sos1 contacts, whereas the role of Grb2 in simultaneously binding LAT and Sos1 is not recapitulated by the model. The weak phosphorylation dependence of Grb2 interactions likely reflects the resolution of the model, which captures chemically specific residue–residue interactions but not site-specific recognition between short linear motifs and folded binding domains [83].

## Conclusions

We combined the CALVADOS model for IDRs and multi-domain proteins with the iSoLFv2 lipid model by introducing protein–lipid cross-interactions based on the potentials and combination rules of the original models. After calibrating a key protein–lipid interaction parameter against transmembrane protein insertion and orientation, we validated the model against experimental transfer free energies for WLXLL pentapeptides. The model reproduces the trends in transfer free-energy in good agreement with both Martini 2 and experiments, although it overestimates the energetic separation between polar and hydrophobic residues.

Through direct comparison with an experimentally refined ensemble obtained from Martini simulations, we showed that the model accurately predicts the conformational properties of the human growth hormone receptor. These include not only the global dimensions of the disordered ICD, but also the orientations of the TMD and ECD, and residue-resolved protein–lipid contacts. Although this validation is limited to a single receptor, the results suggest that MEM-CALVADOS can provide accurate conformational ensembles of membrane proteins containing long IDRs and may offer an alternative to approaches that infer membrane-bound conformations from molecular modeling of isolated disordered regions.

Finally, application of the model to LAT, Grb2, and Sos1 showed that it recapitulates features of the phosphorylation-dependent interaction network, including enhanced recruitment of Sos1 to the membrane interface and the involvement of the SH2 domain of Grb2 in the interactions with LAT. However, the role of Grb2 in simultaneously binding LAT and Sos1 is not fully captured, indicating that site-specific interactions between known binding sites and motifs may be required for a more complete description of multivalent signaling assemblies.

Overall, MEM-CALVADOS provides a computationally efficient model for investigating flexible trans-membrane proteins and protein assembly at bilayer–water interfaces. As such, we envision the model to extend to IDR-containing membrane proteins the range of applications of CALVADOS and similar residue-level models [30, 33, 35, 36, 38, 40], including proteome-wide predictions of conformational properties [29,32, development of machine-learning models from simulation data [31, 84, 85], and protein design [86–88]. An important limitation of the residue-level resolution of the model is the neglect of site-specific interactions. As recognition between folded domains and short linear motifs often organizes multivalent interaction networks of signaling proteins [8], introducing additional interactions between known binding sites and motifs into the model could improve the description of such assemblies [89]. Further, extending the model to include additional lipid species, cholesterol, and capture asymmetric bilayers could enable computational studies of the coupling between protein assembly and lateral membrane organization [90] and of how membrane composition regulates membrane-associated condensate formation and transmembrane coupling [91, 92].

## Code and Data Availability

The implementation of MEM-CALVADOS v1 presented in this work is available as a branch of the CAL-VADOS package (https://github.com/KULL-Centre/CALVADOS/tree/mem-calvados) as well as at https://github.com/gitesei/MEM-CALVADOS. Scripts and data to reproduce the analyses presented in this work are available at https://github.com/gitesei/_2026_MEM-CALVADOS.

## Supporting Information

The Supporting Information includes finite-size effects on membrane properties; convergence of membrane properties; parameter scans for fine-tuning lipid parameters; comparisons of predicted transmembrane-segment assignments and TMD tilt angles with reference data; convergence of conformational properties of the growth hormone receptor; convergence of properties of the LAT–Grb2–Sos1 system; and the Grb2– Sos1 contact profile.

## Acknowledgments

We thank K. Lindorff-Larsen for comments and suggestions. This work was supported by the Swedish Research Council through grant agreement no. 2024-04539. Simulations were enabled by resources provided by LUNARC, The Centre for Scientific and Technical Computing at Lund University, and by the National Academic Infrastructure for Supercomputing in Sweden (NAISS), partially funded by the Swedish Research Council through grant agreement no. 2022-06725. The authors acknowledge a donation to Water Science Lab from Sandberg Development.

