## Supporting Information for "MEM-CALVADOS: A Residue-Level Model for Flexible Proteins at Membrane Interfaces"

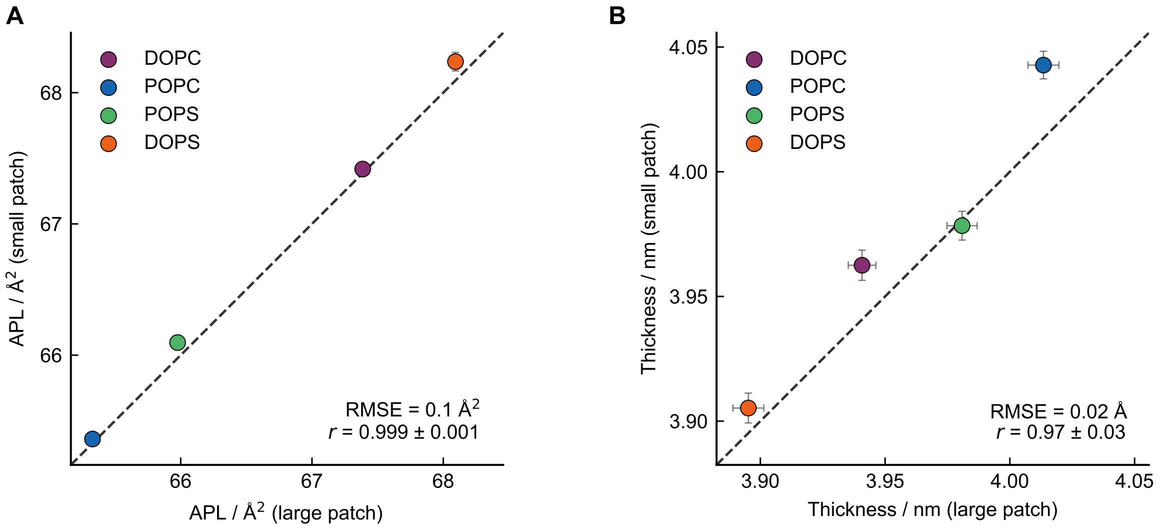


Figure S1: Finite-size effects on membrane properties. (A) Area per lipid (APL) and (B) phosphate–phosphate bilayer thickness from simulations of lipid bilayer patches of different sizes (large patch: 25 nm×25 nm; small patch: 12 nm×12 nm). Data are shown for DOPC, POPC, POPS, and DOPS bilayers at 297.15 K. Error bars are standard errors estimated by blocking analysis.

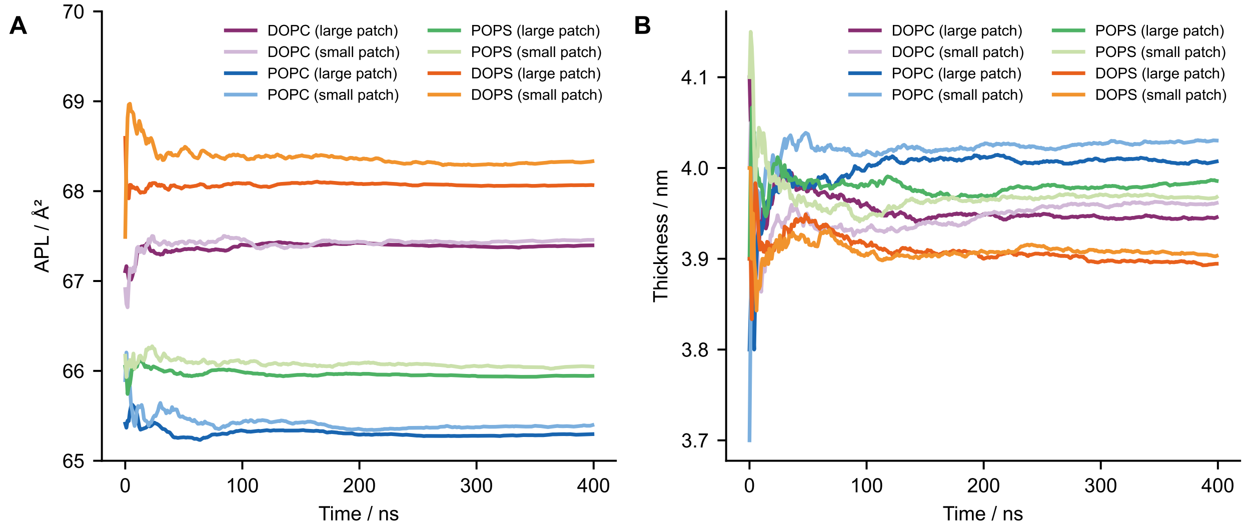


Figure S2: Convergence of membrane properties. Cumulative averages of the (A) area per lipid (APL) and (B) phosphate–phosphate bilayer thickness from simulations of lipid bilayer patches of different sizes (large patch: 25 nm×25 nm; small patch: 12 nm×12 nm). Data are shown for DOPC, POPC, POPS, and DOPS bilayers at 297.15 K.

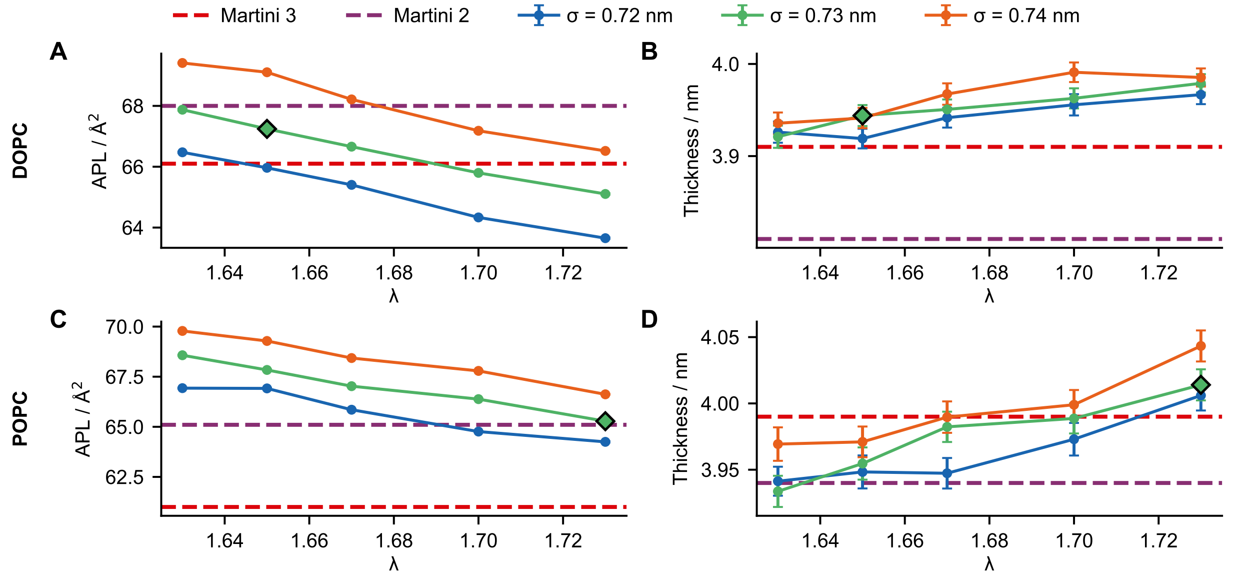


Figure S3: Fine-tuning of the tail bead parameters. (A, C) Area per lipid (APL) and (B, D) phosphate–phosphate bilayer thickness as functions of the tail bead $\lambda$ value, for different values of $\sigma$. Data are shown for (A, B) DOPC and (C, D) POPC bilayers at 297.15 K. Horizontal dashed lines indicate the corresponding reference values from Martini 3 (red) and Martini 2 (purple) [1]. Diamonds indicate the selected optimal parameters. Error bars are standard errors estimated by blocking analysis.

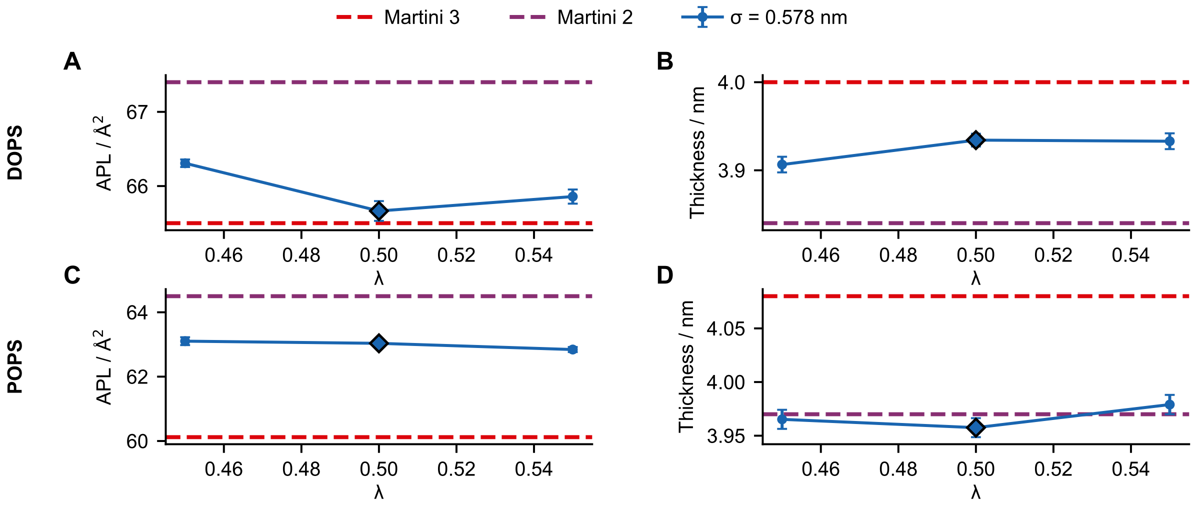


Figure S4: Fine-tuning of the Pho bead $\lambda$ value. (A, C) Area per lipid (APL) and (B, D) phosphate–phosphate bilayer thickness as functions of the Pho bead $\lambda$ value for a fixed Pho bead $\sigma$ of 0.578 nm. Data are shown for (A, B) DOPS and (C, D) POPS bilayers at 297.15 K. Horizontal dashed lines indicate the corresponding reference values from Martini 3 (red) and Martini 2 (purple) [1]. Diamonds indicate the selected $\lambda$ value. Error bars are standard errors estimated by blocking analysis.

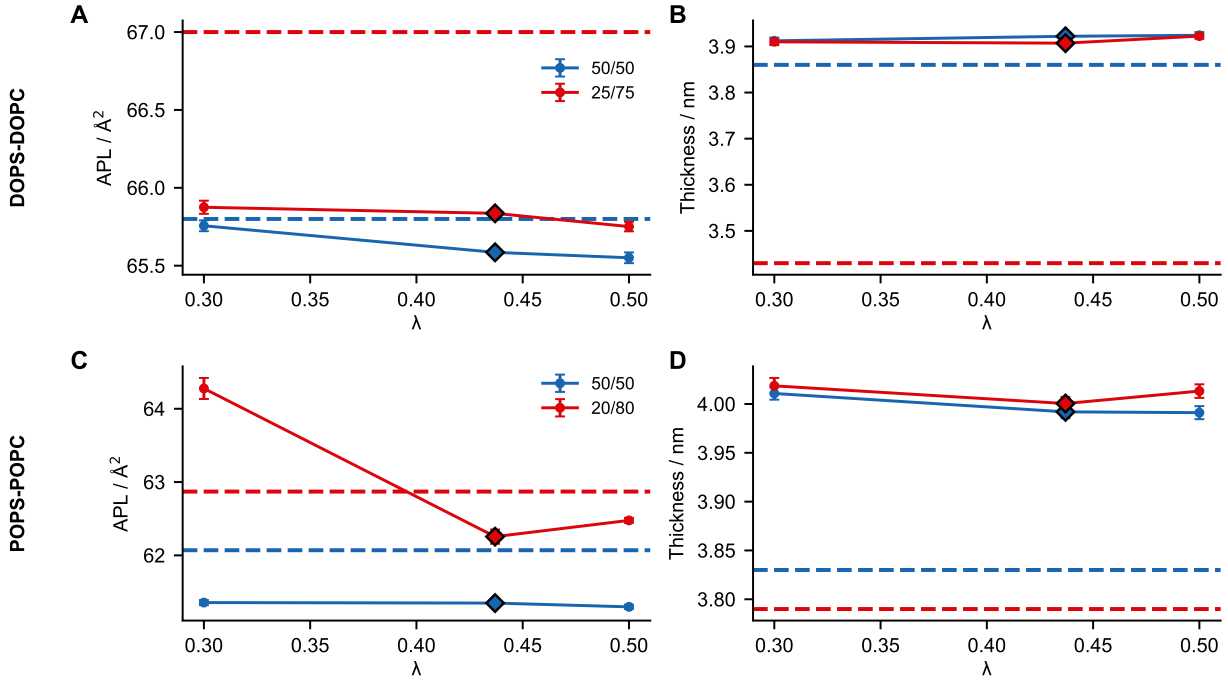


Figure S5: Fine-tuning of the Cho bead $\lambda$ value. (A, C) Area per lipid (APL) and (B, D) phosphate–phosphate bilayer thickness as functions of the Cho bead $\lambda$ value for a fixed Cho bead $\sigma$ of 0.650 nm. Data are shown for mixed (A, B) DOPS/DOPC and (C, D) POPS/POPC bilayers at different molar ratios. Horizontal dashed lines indicate the reference from all-atom simulations from Veretenenko et al. for DOPS/DOPC and from Melcr et al. for POPS/POPC [2–4]. MEM-CALVADOS simulations were performed at the temperature used in the corresponding reference study. Diamonds indicate the selected parameters. Error bars are standard errors estimated by blocking analysis.

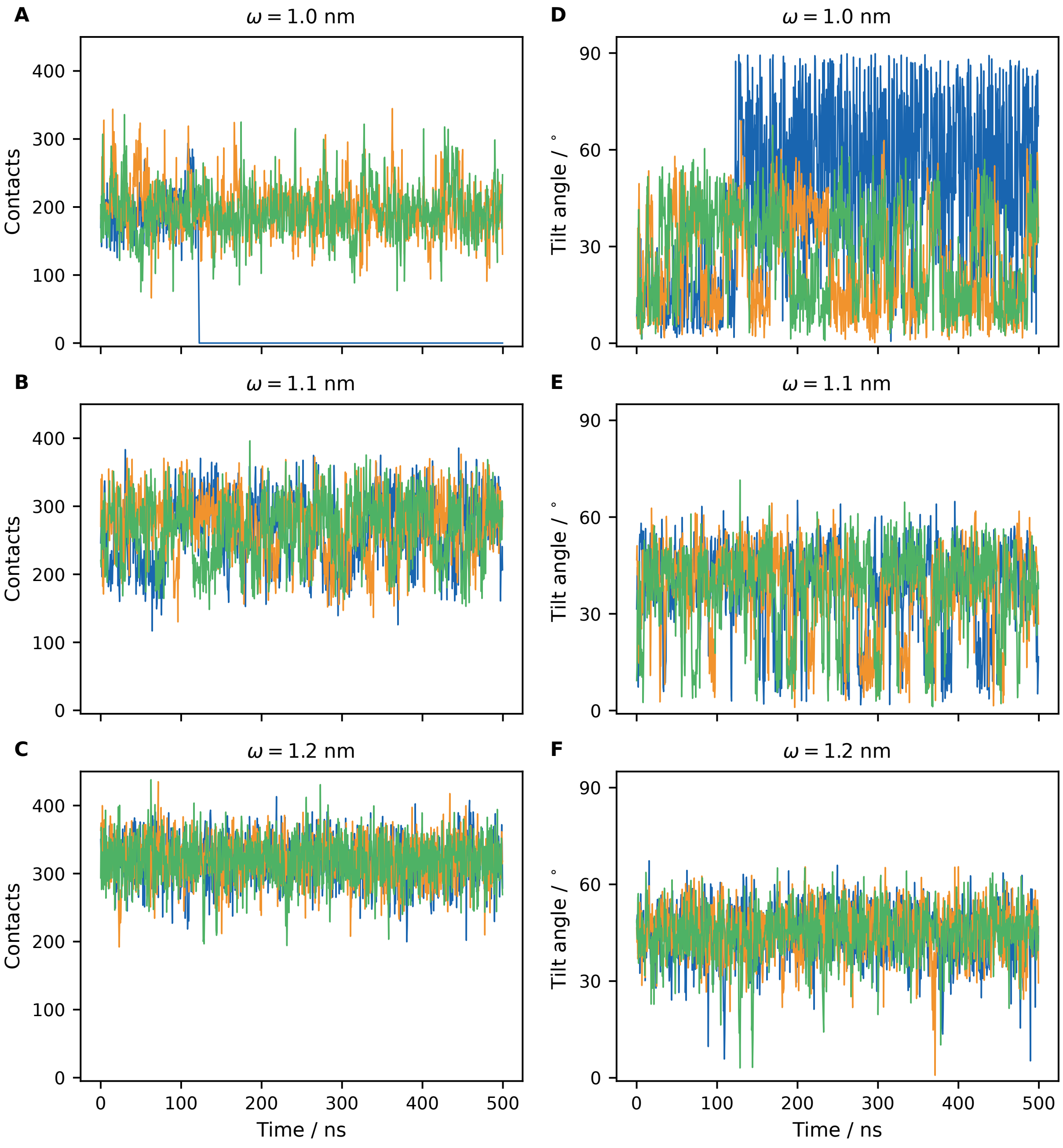


Figure S6: Time series of (A–C) the total number of protein–tail contacts and (D–F) the TMD tilt angle for the helix hairpin of $F_1F_o$ ATP synthase subunit c (PDB 1A91) simulated in three independent replicas (blue, orange, and green) using (A, D) $\omega = 1.0$ nm, (B, E) $\omega = 1.1$ nm, and (C, F) $\omega = 1.2$ nm.

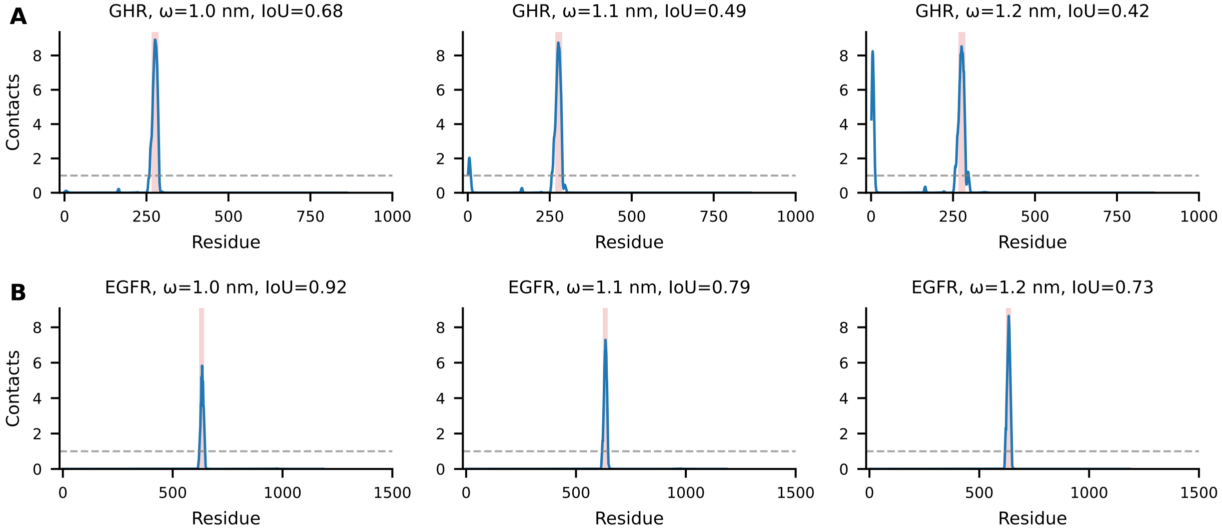


Figure S7: Comparison between transmembrane segments predicted by DeepTMHMM (red shading) and per-residue contacts between protein and lipid tail beads obtained from MEM-CALVADOS simulations (blue curves) for (A) GHR and (B) EGFR at different $\omega$ values. The horizontal dashed lines indicate the threshold of one contact used to convert the contact profiles into transmembrane-segment assignments.

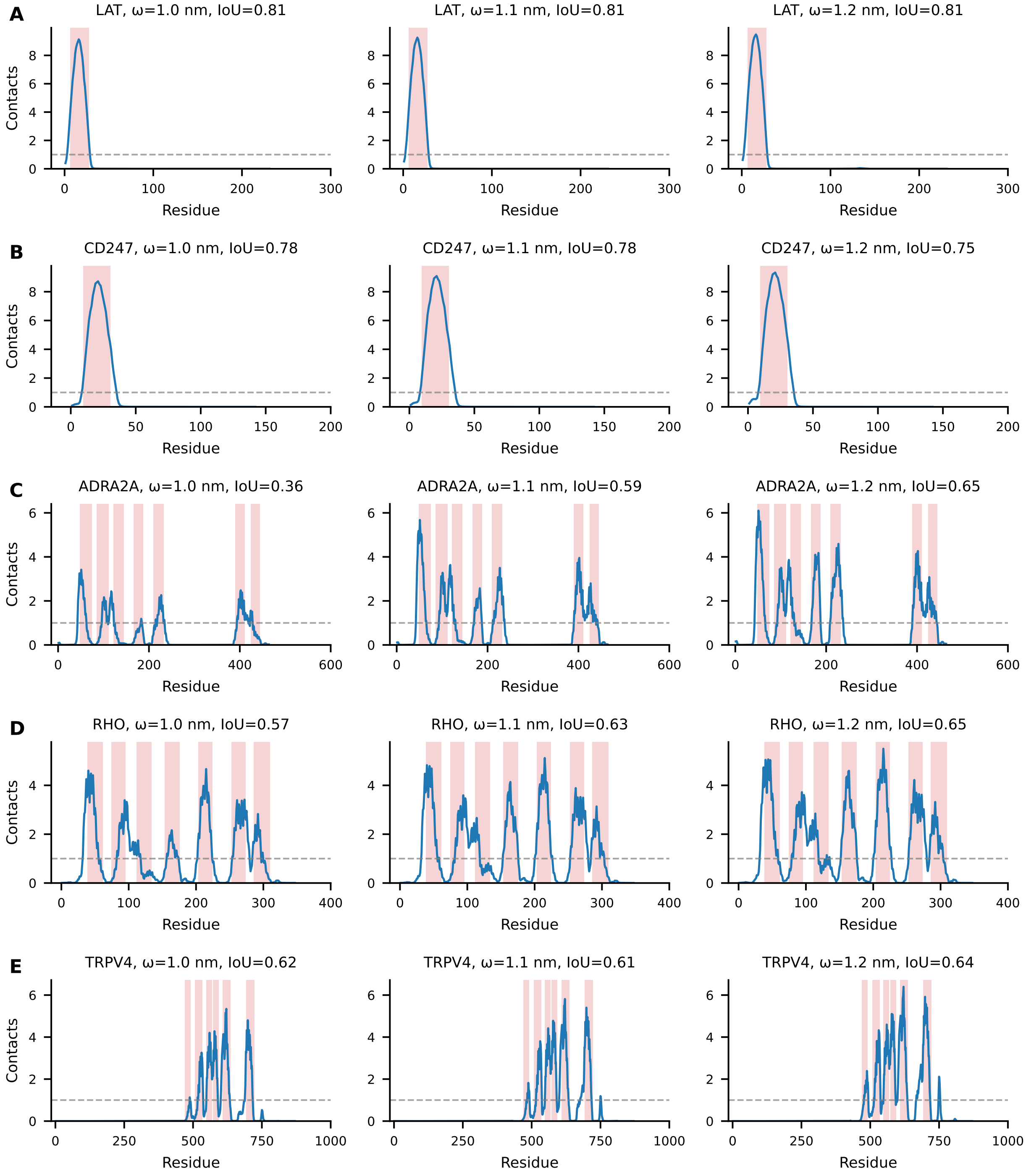


Figure S8: Comparison between transmembrane segments predicted by DeepTMHMM (red shading) and per-residue contacts between protein and lipid tail beads obtained from MEM-CALVADOS simulations (blue curves) for (A) LAT, (B) CD247, (C) ADRA2A, (D) RHO, and (E) TRPV4 at different $\omega$ values. The horizontal dashed lines indicate the threshold of one contact used to convert the contact profiles into transmembrane-segment assignments.

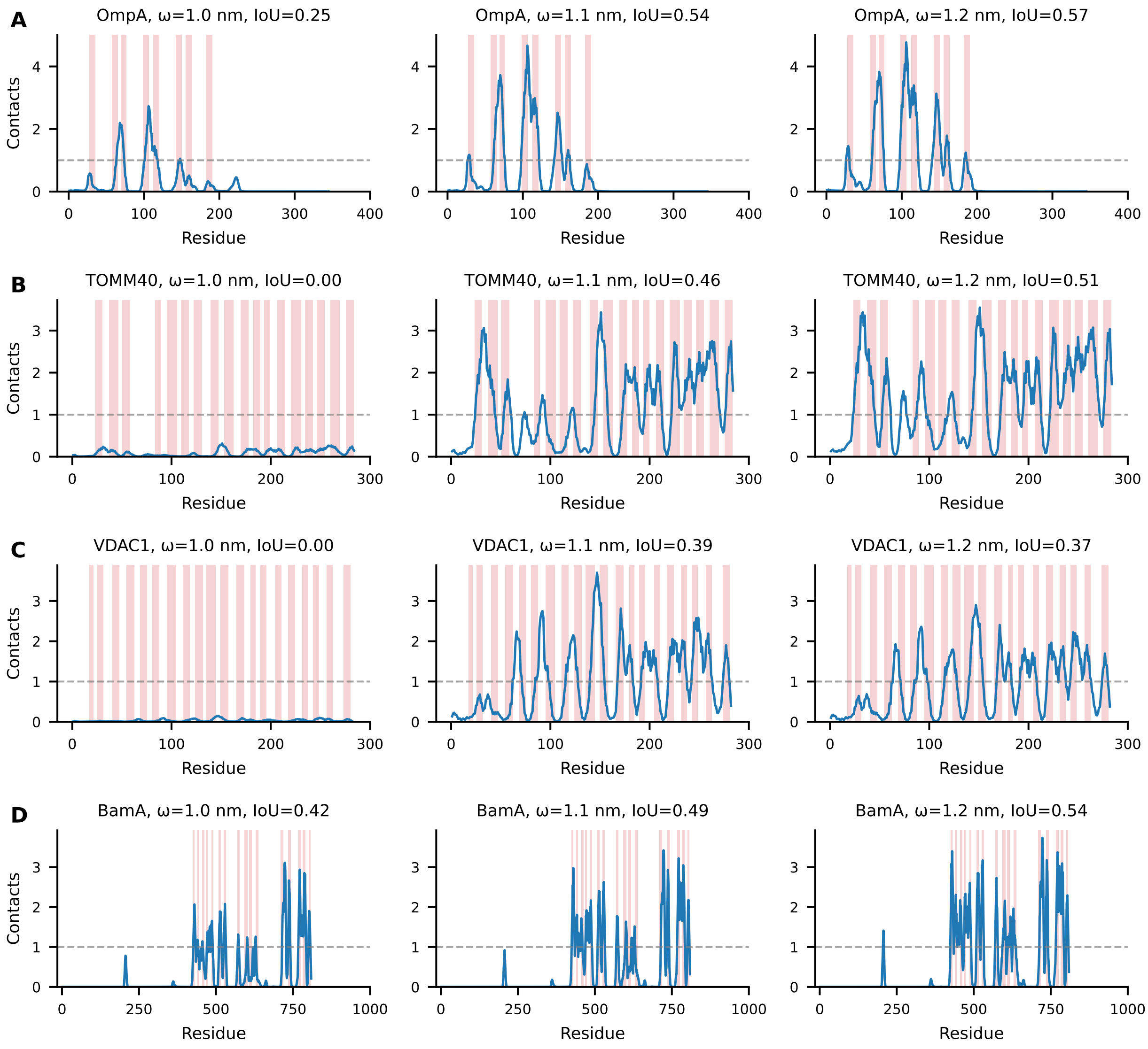


Figure S9: Comparison between transmembrane segments predicted by DeepTMHMM (red shading) and per-residue contacts between protein and lipid tail beads obtained from MEM-CALVADOS simulations (blue curves) for (A) OmpA, (B) TOMM40, (C) VDAC1, and (D) BamA at different $\omega$ values. The horizontal dashed lines indicate the threshold of one contact used to convert the contact profiles into transmembrane-segment assignments.

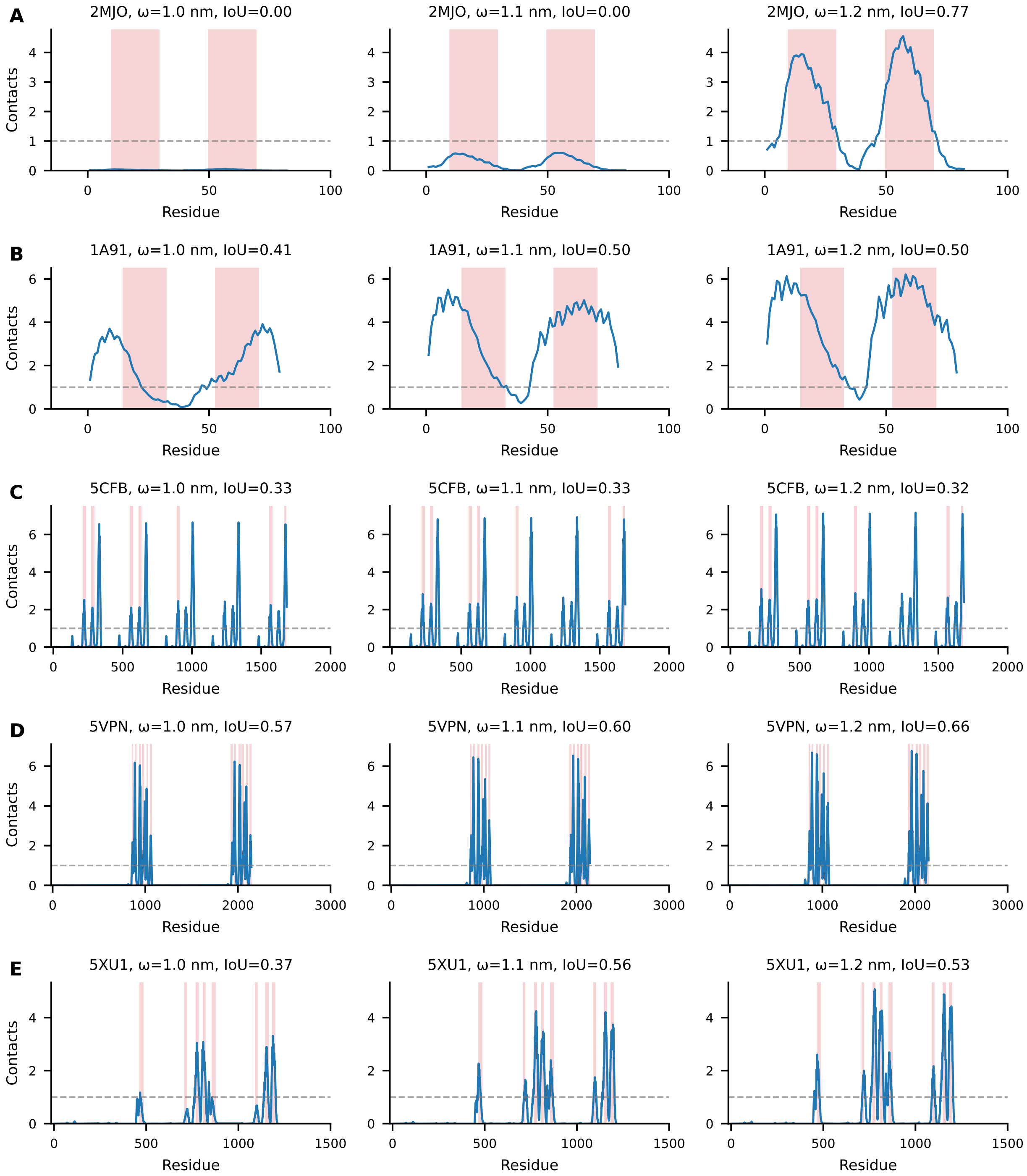


Figure S10: Comparison between transmembrane segments predicted by DeepTMHMM (red shading) and per-residue contacts between protein and lipid tail beads obtained from MEM-CALVADOS simulations (blue curves) for (A) 2MJO, (B) 1A91, (C) 5CFB, (D) 5VPN, and (E) 5XU1 at different $\omega$ values. The horizontal dashed lines indicate the threshold of one contact used to convert the contact profiles into transmembrane-segment assignments.

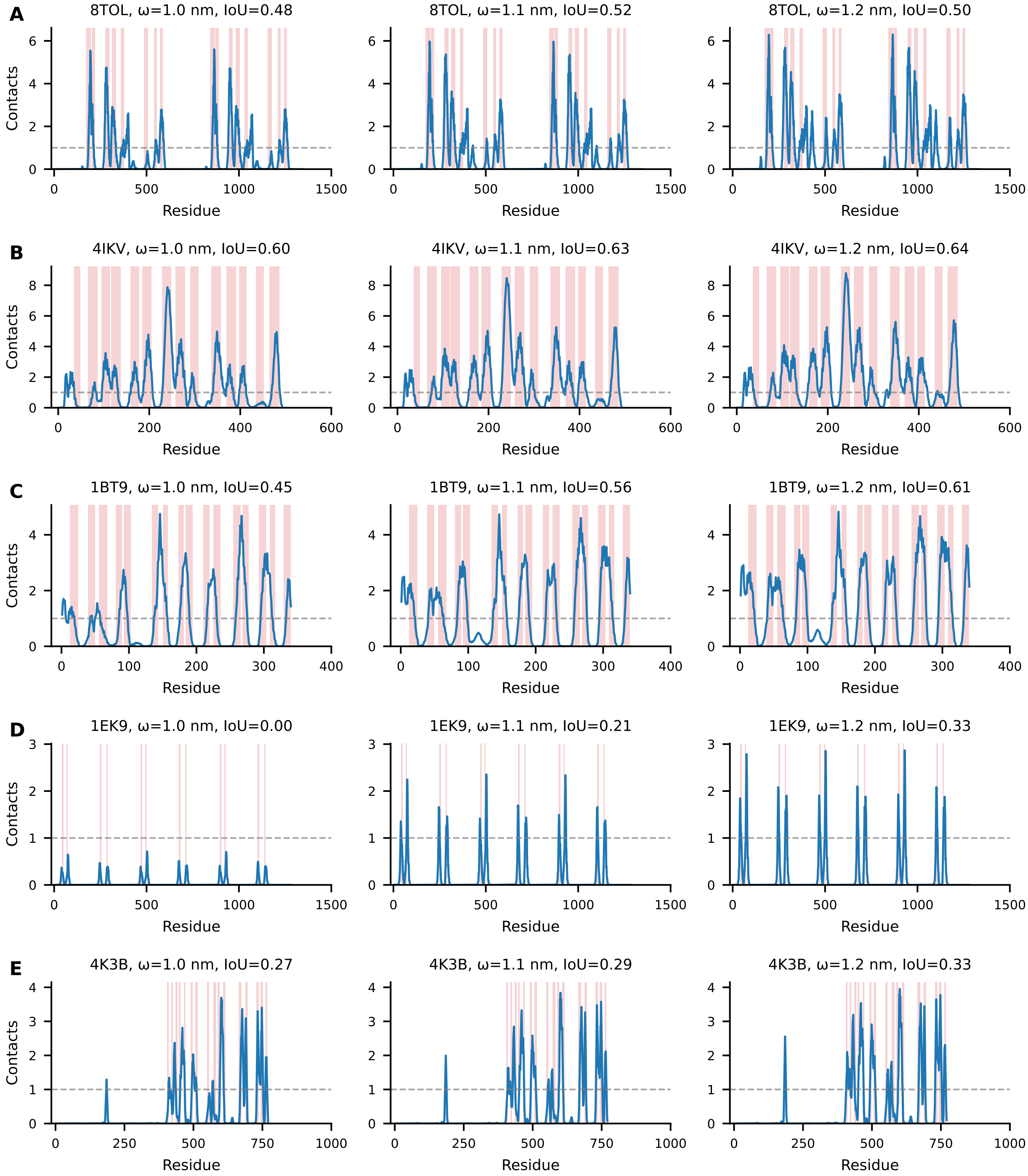


Figure S11: Comparison between transmembrane segments predicted by DeepTMHMM (red shading) and per-residue contacts between protein and lipid tail beads obtained from MEM-CALVADOS simulations (blue curves) for (A) 8TOL, (B) 4IKV, (C) 1BT9, (D) 1EK9, and (E) 4K3B at different $\omega$ values. The horizontal dashed lines indicate the threshold of one contact used to convert the contact profiles into transmembrane-segment assignments.

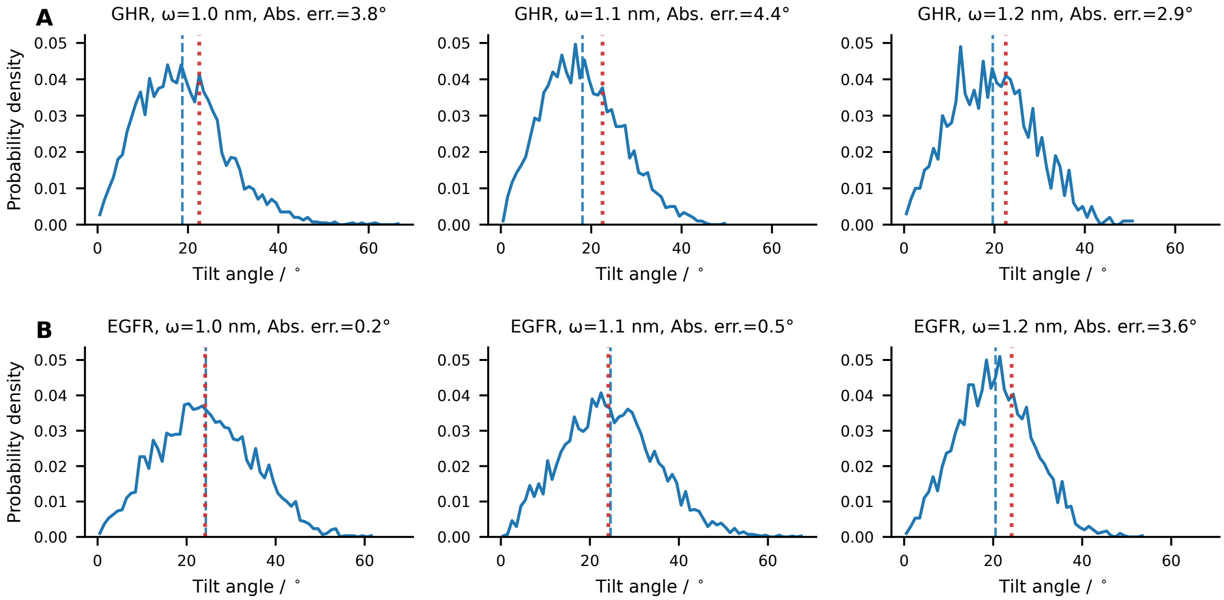


Figure S12: Comparison between the TMD tilt angle relative to the membrane normal predicted by MemPrO (red dotted line) and the probability density of the TMD tilt angle obtained from MEM-CALVADOS simulations (blue curve) for (A) GHR and (B) EGFR at different $\omega$ values. The blue dashed line indicates the mean of the distribution, and the corresponding absolute error relative to the MemPrO reference is reported in each panel.

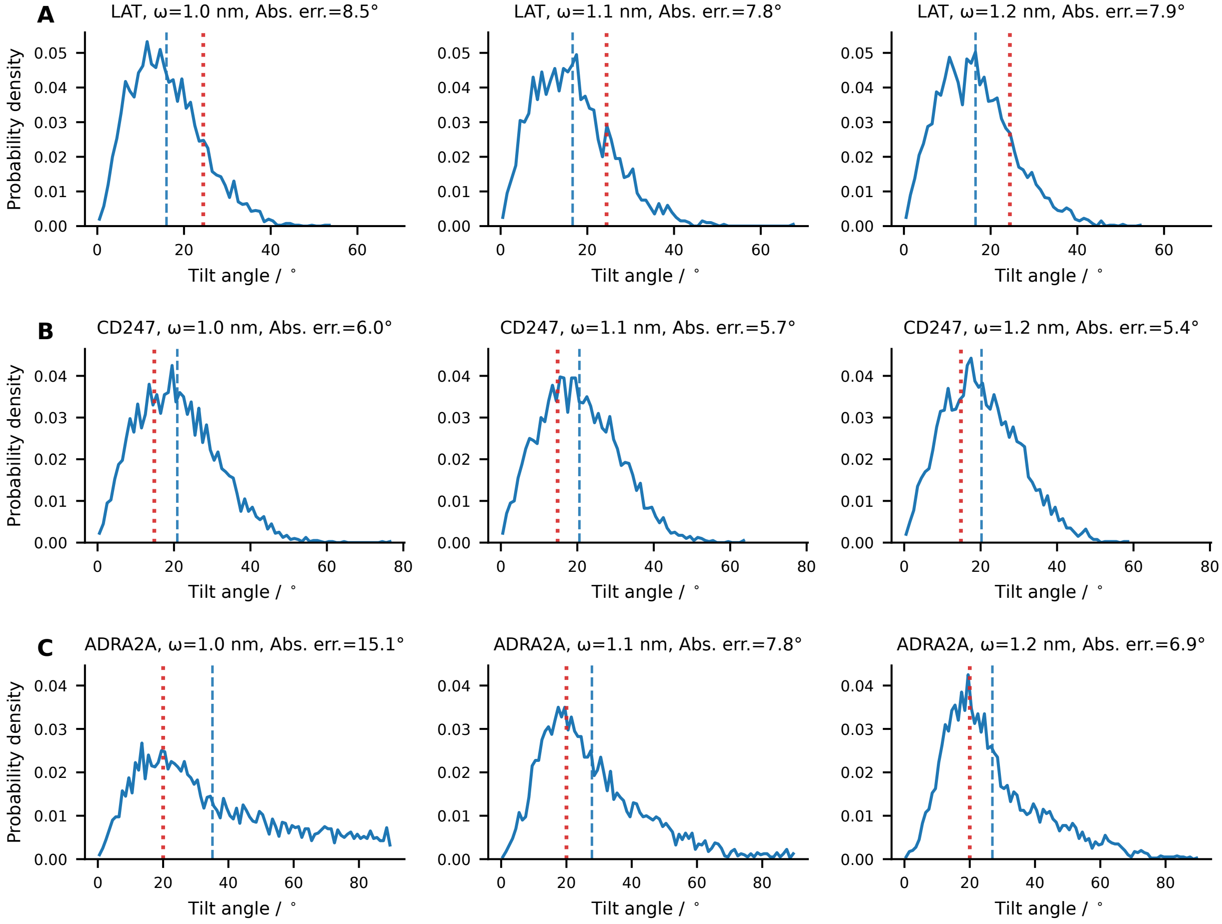


Figure S13: Comparison between the TMD tilt angle relative to the membrane normal predicted by MemPrO (red dotted line) and the probability density of the TMD tilt angle obtained from MEM-CALVADOS simulations (blue curve) for (A) LAT, (B) CD247, and (C) ADRA2A at different $\omega$ values. The blue dashed line indicates the mean of the distribution, and the corresponding absolute error relative to the MemPrO reference is reported in each panel.

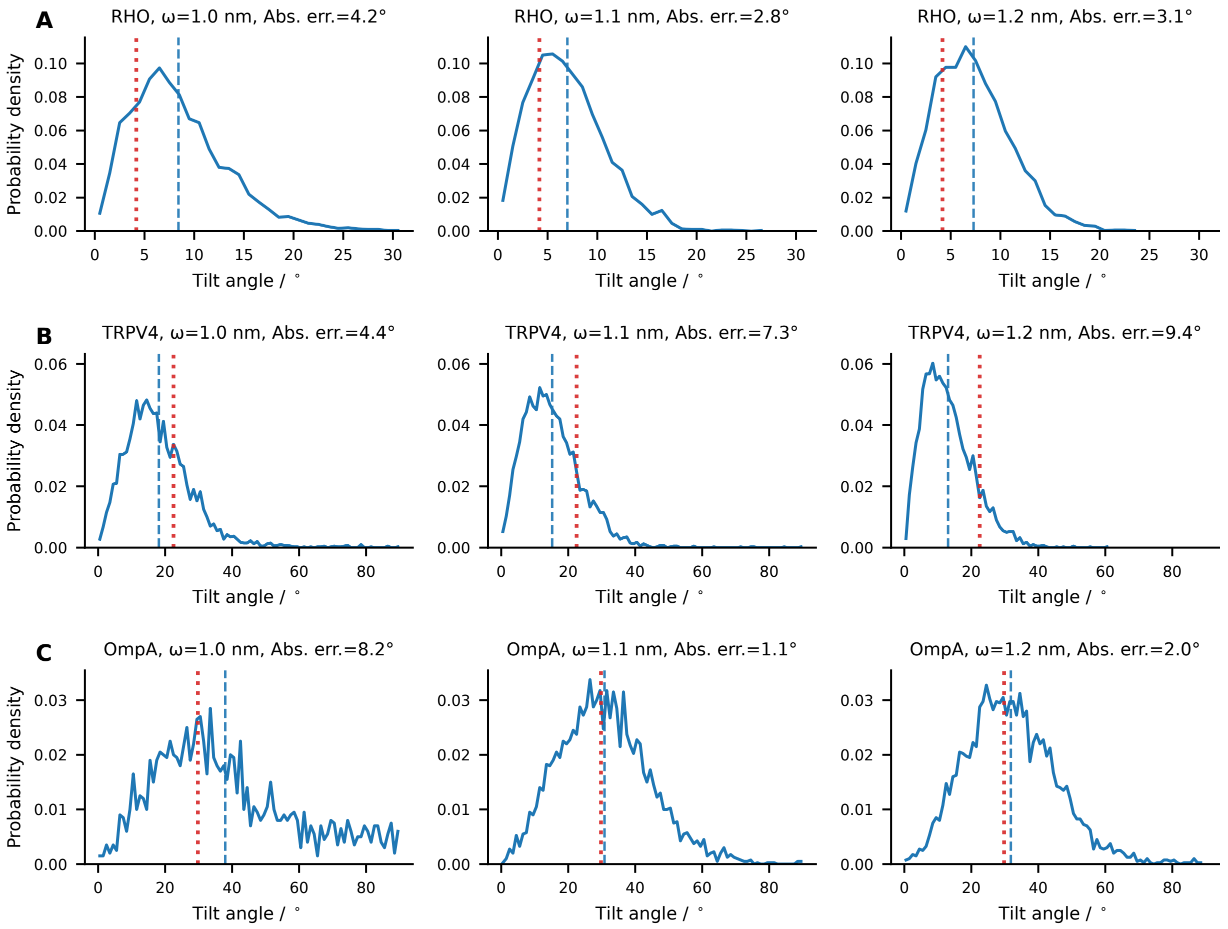


Figure S14: Comparison between the TMD tilt angle relative to the membrane normal predicted by MemPrO (red dotted line) and the probability density of the TMD tilt angle obtained from MEM-CALVADOS simulations (blue curve) for (A) RHO, (B) TRPV4, and (C) OmpA at different $\omega$ values. The blue dashed line indicates the mean of the distribution, and the corresponding absolute error relative to the MemPrO reference is reported in each panel.

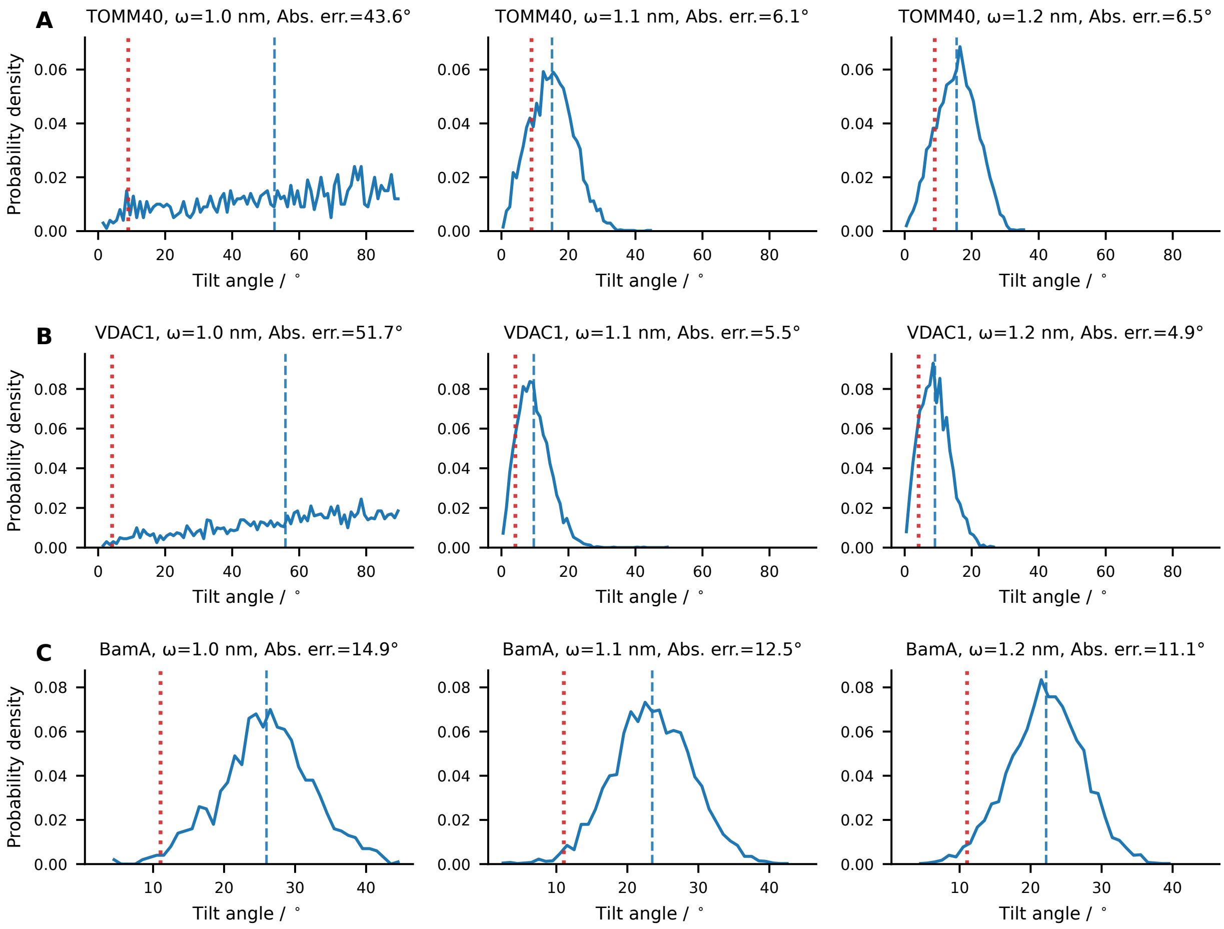


Figure S15: Comparison between the TMD tilt angle relative to the membrane normal predicted by MemPrO (red dotted line) and the probability density of the TMD tilt angle obtained from MEM-CALVADOS simulations (blue curve) for (A) TOMM40, (B) VDAC1 and (C) BamA at different $\omega$ values. The blue dashed line indicates the mean of the distribution, and the corresponding absolute error relative to the MemPrO reference is reported in each panel.

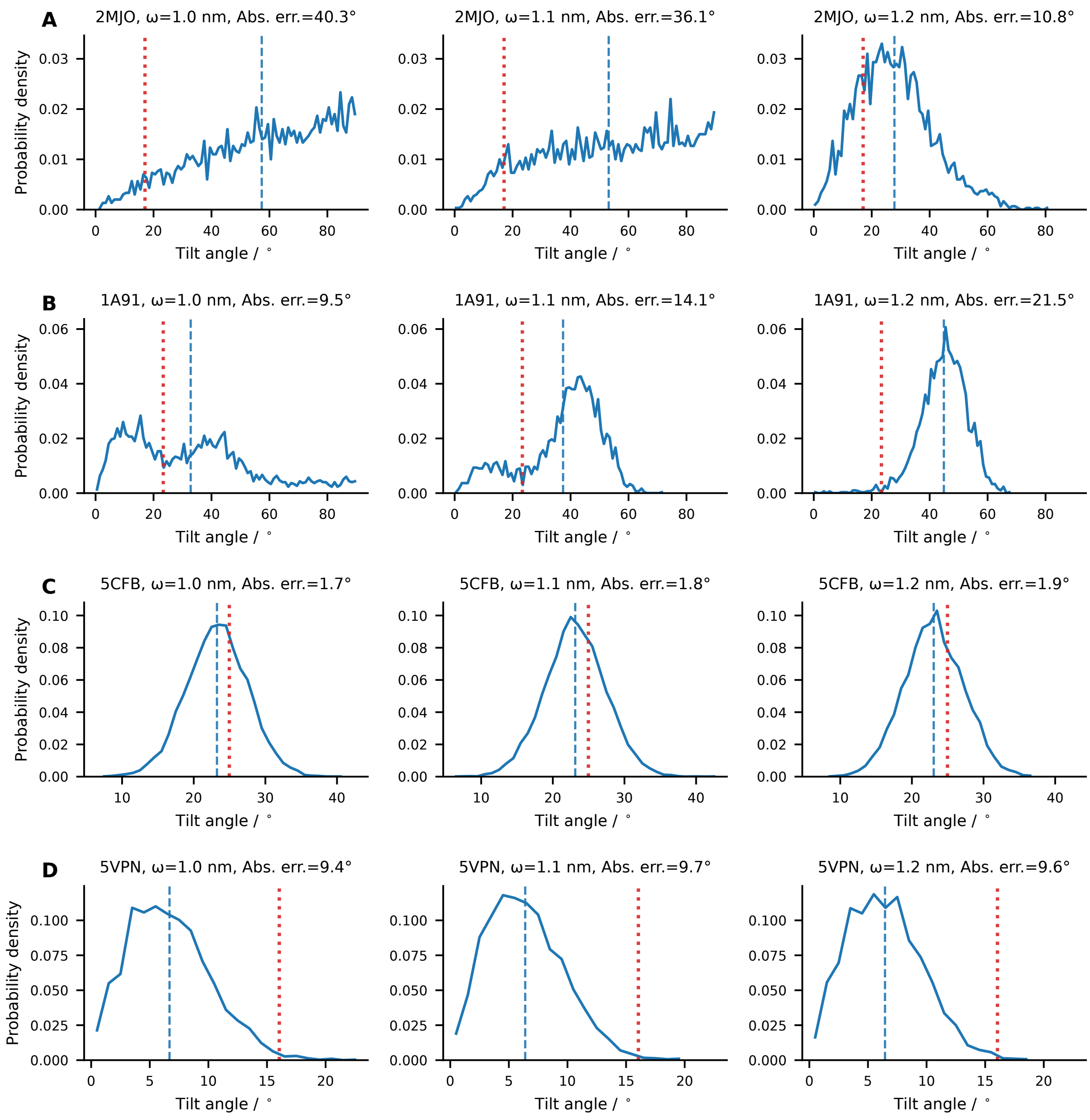


Figure S16: Comparison between the TMD tilt angle relative to the membrane normal predicted by MemPrO (red dotted line) and the probability density of the TMD tilt angle obtained from MEM-CALVADOS simulations (blue curve) for (A) 2MJO, (B) 1A91, (C) 5CFB, and (D) 5VPN at different $\omega$ values. The blue dashed line indicates the mean of the distribution, and the corresponding absolute error relative to the MemPrO reference is reported in each panel.

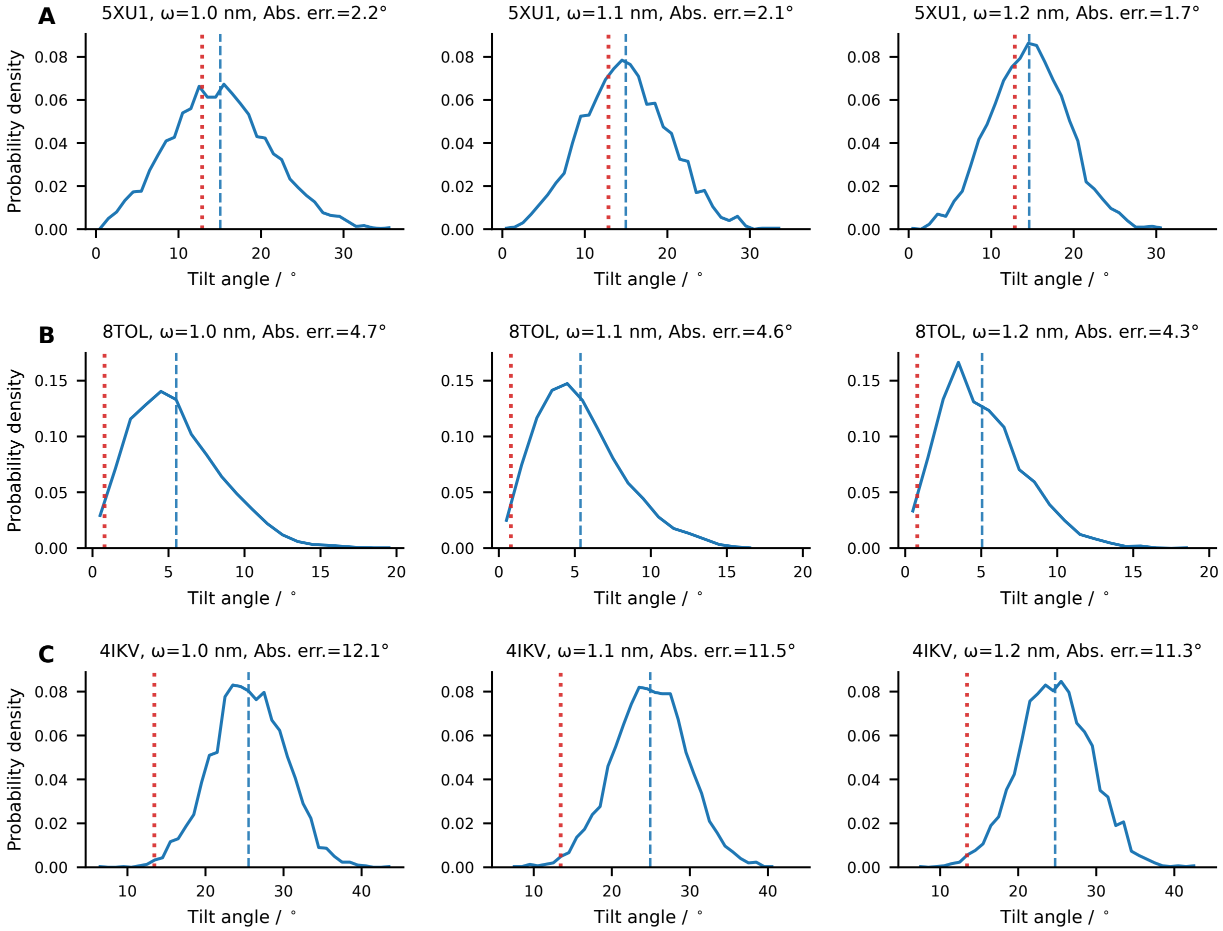


Figure S17: Comparison between the TMD tilt angle relative to the membrane normal predicted by MemPrO (red dotted line) and the probability density of the TMD tilt angle obtained from MEM-CALVADOS simulations (blue curve) for (A) 5XU1, (B) 8TOL, and (C) 4IKV at different $\omega$ values. The blue dashed line indicates the mean of the distribution, and the corresponding absolute error relative to the MemPrO reference is reported in each panel.

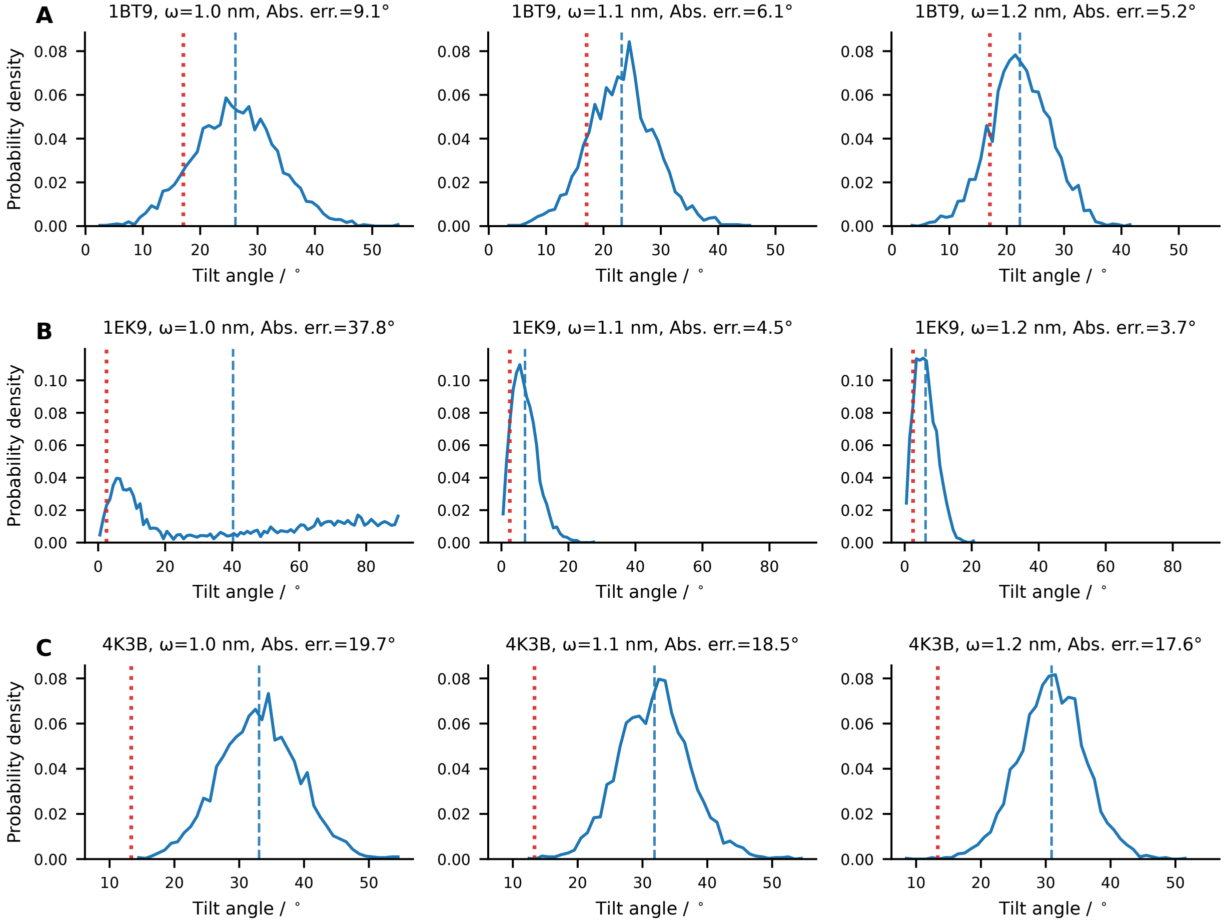


Figure S18: Comparison between the TMD tilt angle relative to the membrane normal predicted by MemPrO (red dotted line) and the probability density of the TMD tilt angle obtained from MEM-CALVADOS simulations (blue curve) for (A) 1BT9, (B) 1EK9, and (C) 4K3B at different $\omega$ values. The blue dashed line indicates the mean of the distribution, and the corresponding absolute error relative to the MemPrO reference is reported in each panel.

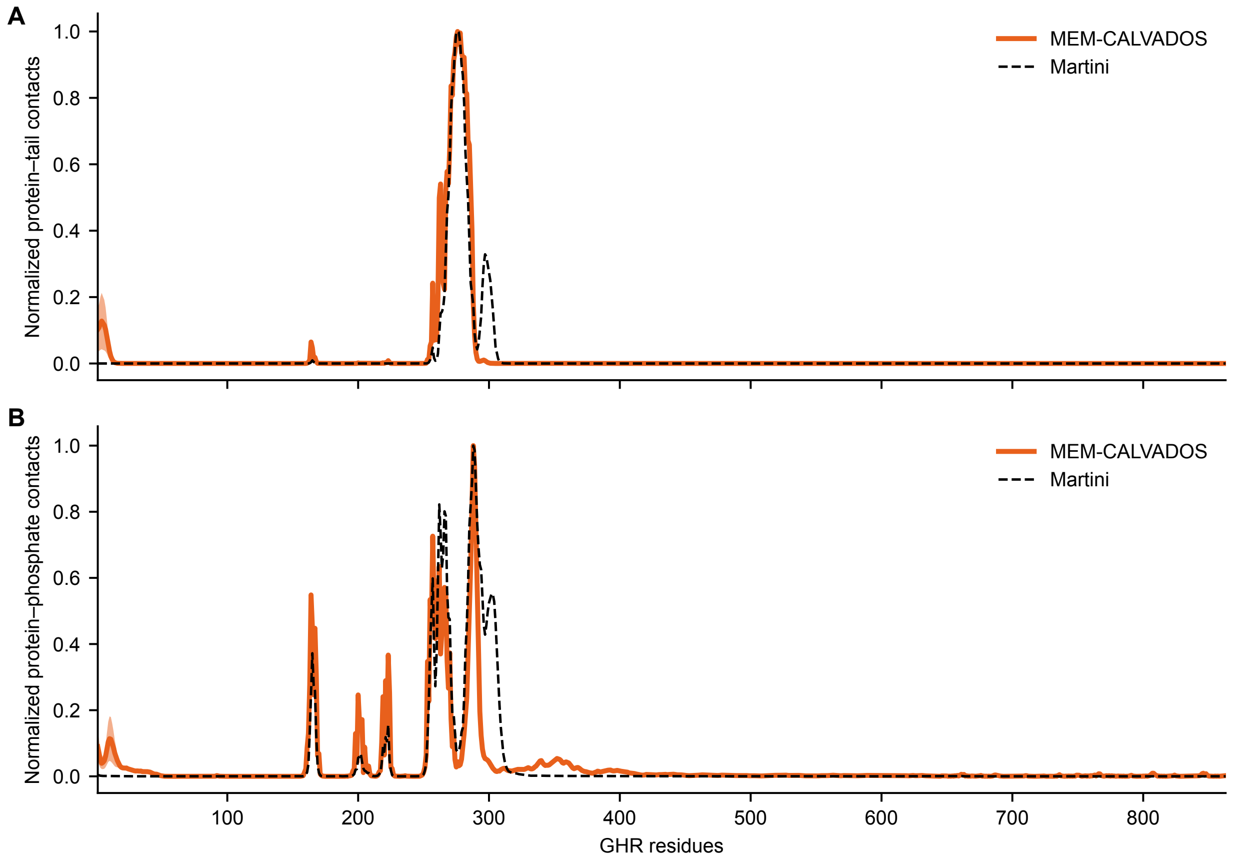


Figure S19: (A) Protein–lipid tail contacts and (B) protein–phosphate contacts, normalized by the maximum value, for MEM-CALVADOS (orange) and the Martini simulations from Kassem et al. (black) [5]. Martini contacts were calculated using the same analysis procedure as for MEM-CALVADOS, with contacts evaluated between protein BB beads and all lipid tail beads (C1A, D2A, C3A, C4A, C1B, C2B, C3B, and C4B) or PO4 beads, using the combined production trajectory archived on Zenodo [6].

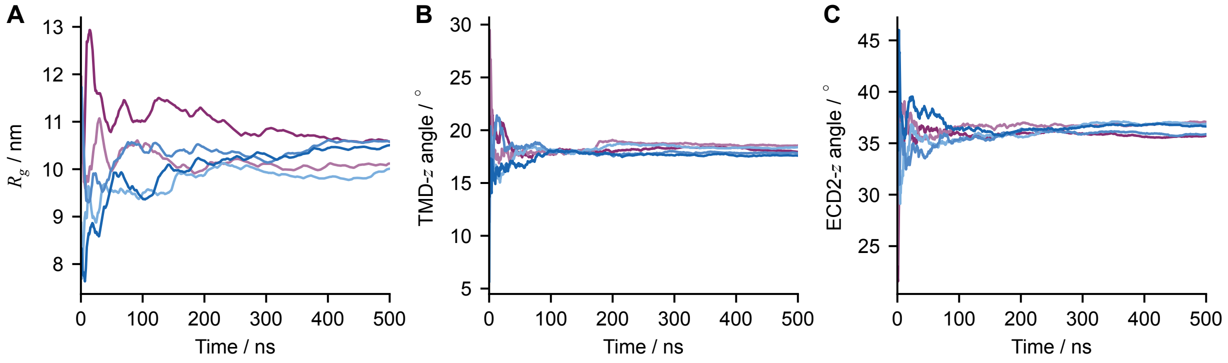


Figure S20: Cumulative averages from five independent simulations of full-length GHR in a POPC bilayer for (A) the radius of gyration, $R_g$, of the full-length protein, (B) the tilt angle of the transmembrane domain, and (C) the angle between the principal axis of the extracellular D2 domain and the bilayer normal.

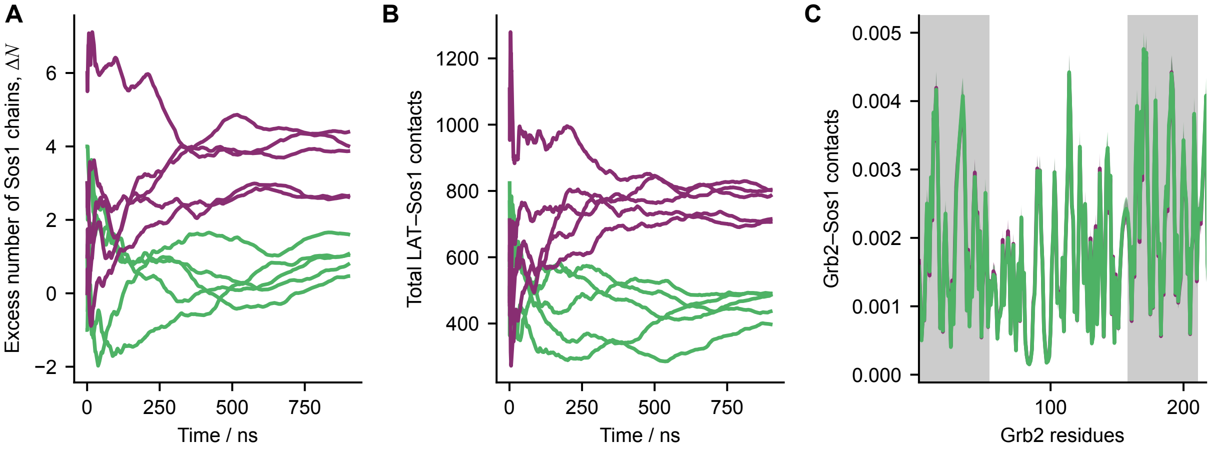


Figure S21: (A) Cumulative averages of the excess number, $\Delta N$, of Sos1 chains in the intracellular region, $-20$ nm $< z_{\mathrm{COM}} < 0$ nm, relative to the corresponding extracellular region, 0 nm $< z_{\mathrm{COM}} <$ 20 nm from five replicas of the LAT–Grb2–Sos1 system with phosphorylated (purple) and unphosphorylated (green) LAT. (B) Cumulative averages of the total number of contacts between LAT and Sos1 from five replicas of the LAT-Grb2-Sos1 system with phosphorylated (purple) and unphosphorylated (green) LAT. (C) Grb2 contacts with Sos1 in the systems with phosphorylated (purple) and unphosphorylated (green) LAT. Solid lines in panel C are means over five independent replicas, and gray shaded areas indicate the SH3 domains.

Table S1: Bonded parameters for the lipid model. The bond and angle parameters in each row apply to bonds and angle triplets starting from the corresponding bead type. TPO and TDO are palmitoyl-oleoyl and dioleoyl tail beads.

| Type | $r_0$ / nm | $k_b$ / kJ mol$^{-1}$ nm$^{-2}$ | $\theta_0$ / rad | $k_\theta$ / kJ mol$^{-1}$ rad$^{-2}$ |
|---|---|---|---|---|
| Ser | 0.442 | 17 660 | 2.240 | 16.000 |
| Cho | 0.402 | 4010 | 2.460 | 7.400 |
| Pho | 0.471 | 3788 | 3.125 | 13.000 |
| Mid | 0.465 | 4734 | 3.125 | 19.600 |
| TPO | 0.530 | 2312 | 3.125 | 9.800 |
| TDO | 0.520 | 2634 | 3.125 | 9.000 |

Table S2: Lipid-bead and amino-acid parameters used for nonbonded interactions. TPO and TDO are palmitoyl-oleoyl and dioleoyl tail beads.

| Bead/residue | $q$ / $e$ | $\lambda$ | $\sigma$ / nm | $\omega$ / nm |
|---|---|---|---|---|
| *Lipid beads* | | | | |
| Ser | 0.000 | 0.447 | 0.588 | – |
| Cho | 1.000 | 0.437 | 0.650 | – |
| Pho | −1.000 | 0.500 | 0.578 | – |
| Mid | 0.000 | 0.469 | 0.507 | – |
| TPO | 0.000 | 1.730 | 0.720 | 1.200 |
| TDO | 0.000 | 1.650 | 0.730 | 1.200 |
| *Amino acids* | | | | |
| Glu | −1.000 | 0.000 | 0.592 | – |
| Asp | −1.000 | 0.093 | 0.558 | – |
| Lys | 1.000 | 0.138 | 0.636 | – |
| Thr | 0.000 | 0.267 | 0.562 | – |
| Val | 0.000 | 0.294 | 0.586 | 1.100 |
| Gln | 0.000 | 0.314 | 0.602 | – |
| Ala | 0.000 | 0.338 | 0.504 | – |
| Pro | 0.000 | 0.347 | 0.556 | – |
| Asn | 0.000 | 0.371 | 0.568 | – |
| His | 0.000 | 0.409 | 0.608 | – |
| Ser | 0.000 | 0.447 | 0.518 | – |
| Ile | 0.000 | 0.513 | 0.618 | 1.100 |
| Met | 0.000 | 0.517 | 0.618 | 1.100 |
| Leu | 0.000 | 0.555 | 0.618 | 1.100 |
| Cys | 0.000 | 0.592 | 0.548 | – |
| Arg | 1.000 | 0.741 | 0.656 | – |
| Gly | 0.000 | 0.754 | 0.450 | – |
| Phe | 0.000 | 0.891 | 0.636 | 1.100 |
| Tyr | 0.000 | 0.951 | 0.646 | 1.100 |
| Trp | 0.000 | 1.033 | 0.678 | 1.100 |

Table S3: Functional forms used for nonbonded interactions between amino-acid or lipid bead types and lipid beads. AH denotes the Ashbaugh–Hatch potential, WCA the Weeks–Chandler–Andersen potential, and SA the stretched attractive potential. TPO and TDO are palmitoyl-oleoyl and dioleoyl tail beads.

| Bead/residue | Lipid Ser | Cho/Pho/Mid | TPO/TDO |
|---|---|---|---|
| *Lipid beads* | | | |
| Ser | AH | AH | WCA |
| Cho | AH | WCA | WCA |
| Pho | AH | WCA | WCA |
| Mid | WCA | WCA | WCA |
| TPO | WCA | WCA | SA |
| TDO | WCA | WCA | SA |
| *Amino acids* | | | |
| Glu | WCA | WCA | WCA |
| Asp | AH | AH | WCA |
| Lys | AH | AH | WCA |
| Thr | AH | AH | WCA |
| Val | WCA | WCA | SA |
| Gln | AH | AH | WCA |
| Ala | AH | AH | WCA |
| Pro | AH | AH | WCA |
| Asn | AH | AH | WCA |
| His | AH | AH | WCA |
| Ser | AH | AH | WCA |
| Ile | WCA | WCA | SA |
| Met | WCA | WCA | SA |
| Leu | WCA | WCA | SA |
| Cys | AH | AH | WCA |
| Arg | AH | AH | WCA |
| Gly | WCA | WCA | WCA |
| Phe | WCA | WCA | SA |
| Tyr | WCA | WCA | SA |
| Trp | WCA | WCA | SA |

Table S4: Transmembrane protein structures used to optimize the $\omega$ parameter assigned to hydrophobic residues.

| Protein | Organism | Topology | Source |
|---|---|---|---|
| *Full-length models from the literature* | | | |
| Human growth hormone receptor (GHR) | *H. sapiens* | Single-pass $\alpha$-helical | Kassem et al. [5] |
| Epidermal growth factor receptor (EGFR) | *H. sapiens* | Single-pass $\alpha$-helical | Srinivasan et al. [7] |
| *AlphaFold Database models* | | | |
| Linker for activation of T-cells family member 1 (LAT) | *H. sapiens* | Single-pass $\alpha$-helical | AF-O43561-4-F1 |
| T-cell surface glycoprotein CD3 zeta chain (CD247) | *H. sapiens* | Single-pass $\alpha$-helical | AF-P20963-F1 |
| Alpha-2A adrenergic receptor (ADRA2A) | *H. sapiens* | Multipass $\alpha$-helical | AF-P08913-F1 |
| Rhodopsin (RHO) | *H. sapiens* | Multipass $\alpha$-helical | AF-P08100-F1 |
| Transient receptor potential cation channel subfamily V member 4 (TRPV4) | *H. sapiens* | Multipass $\alpha$-helical | AF-Q9HBA0-F1 |
| Outer membrane protein A (OmpA) | *E. coli* | $\beta$-barrel | AF-P0A910-F1 |
| Mitochondrial import receptor subunit TOM40 homolog (TOMM40) | *H. sapiens* | $\beta$-barrel | AF-O96008-F1 |
| Voltage-dependent anion-selective channel protein 1 (VDAC1) | *H. sapiens* | $\beta$-barrel | AF-P21796-F1 |
| Outer membrane protein assembly factor BamA | *E. coli* | $\beta$-barrel | AF-P0A940-F1 |
| *MemPrO structures* | | | |
| p75 neurotrophin receptor transmembrane domain | *R. norvegicus* | Single-pass $\alpha$-helical dimer | PDB 2MJO |
| $F_1F_o$ ATP synthase subunit c | *E. coli* | Multipass $\alpha$-helical | PDB 1A91 |
| Human glycine receptor $\alpha 3$ | *H. sapiens* | Multipass $\alpha$-helical | PDB 5CFB |
| Quinol:fumarate reductase | *E. coli* | Multipass $\alpha$-helical | PDB 5VPN |
| Non-canonical ABC transporter | *S. pneumoniae* | Multipass $\alpha$-helical | PDB 5XU1 |
| nhTMEM16 lipid scramblase | *F. vanettenii* | Multipass $\alpha$-helical | PDB 8TOL |
| Peptide transporter POT | *G. kaustophilus* | Multipass $\alpha$-helical | PDB 4IKV |
| OmpF porin | *E. coli* | $\beta$-barrel | PDB 1BT9 |
| TolC outer membrane protein | *E. coli* | $\beta$-barrel complex | PDB 1EK9 |
| Outer membrane protein assembly factor BamA | *N. gonorrhoeae* | $\beta$-barrel | PDB 4K3B |
